# β-tubulin tyrosination tunes glutamylation and parasite motility in *Trypanosoma brucei*

**DOI:** 10.64898/2026.11.20.767850

**Authors:** Jeremy Vincent, Norma Veltz, Lucien Crobu, Rosa Milagros Corrales, Augustin Tuyishime, Christine Girard-Blanc, Gaël Moneron, Serge Urbach, Yvon Sterkers, Krzysztof Rogowski, Serge Bonnefoy, Philippe Bastin, Maude F Leveque

**Affiliations:** University of Montpellier, CNRS, IRD, MiVEGEC, Montpellier, France; Trypanosome Cell Biology, Institut Pasteur, Université Paris Cité, INSERM U1347, Paris, France; Unit of Signaling and Receptors Dynamics, Institut Pasteur, Université de Paris Cité, CNRS UMR 3571, Paris, France; IGF, Univ. Montpellier, CNRS, INSERM, Montpellier, France; Tubulin Code team, IGH, CNRS, Univ. Montpellier, Montpellier, France

## Abstract

Post-translational modifications of tubulin diversify microtubule function by generating distinct molecular states collectively known as the tubulin code. Whereas the reversible tyrosination cycle is a conserved feature of α-tubulin throughout eukaryotes, in trypanosomatid parasites, β-tubulin also carries a genetically encoded C-terminal tyrosine, which is subjected to detyrosination and whose biological role remains unknown. Here, we show that β-tubulin tyrosination is rare across eukaryotes but highly conserved across trypanosomatids. Using genome editing to uncouple the tyrosination states of α- and β-tubulin in *Trypanosoma brucei*, we demonstrate that the β-tubulin C-terminal tyrosine independently contributes to the regulation of microtubule glutamylation. Rather than depending on α- or β-tubulin alone, glutamylation levels reflect the combined tyrosination states of both subunits. Strikingly, either increasing or reducing β-tubulin tyrosination produces opposing changes in parasite morphometry, single-cell motility and collective migration. Together, these findings establish β-tubulin (de)tyrosination as an evolutionarily conserved component of the trypanosomatid tubulin code that contributes to microtubule regulation and function in *T. brucei*, extending the significance of tubulin tyrosination beyond the canonical α-tubulin tyrosination cycle.

## Introduction

*Trypanosoma brucei* is a flagellated protozoan parasite causing human and animal African trypanosomiasis (sleeping sickness and Nagana, respectively). It belongs to the trypanosomatid lineage (phylum Euglenozoa), which also includes *Trypanosoma cruzi* and *Leishmania* spp., collectively responsible for neglected tropical diseases targeted by the WHO 2021–2030 roadmap [1]. Throughout their life cycles, trypanosomatids rely on a highly organized microtubule (MT) cytoskeleton that defines their cell shape, drives motility and orchestrates essential processes including cell division and morphogenesis [2]. This cytoskeletal network comprises the subpellicular corset, the flagellar axoneme, the mitotic spindle and the microtubule quartet (MtQ), all of which are built from α/β-tubulin heterodimers [3].

Beyond providing structural support, MTs constitute dynamic signalling platforms whose functions depend on the selective recruitment of microtubule-associated proteins (MAPs). This molecular specificity is regulated by the tubulin code, whereby combinations of tubulin isotypes and post-translational modifications (PTMs) generate biochemically distinct MT populations recognized by specific effectors [4]. Unlike most eukaryotes, trypanosomatids express only a limited repertoire of α- and β-tubulin isotypes [3], with *T. brucei* expressing a single α/β-tubulin heterodimer [5]. Consequently, PTMs represent the principal source of MT functional diversity in these parasites.

Among tubulin PTMs, the reversible tyrosination cycle of α-tubulin is one of the best-characterized regulatory systems and has become a paradigm for understanding how the tubulin code controls MT function [4]. The genetically encoded C-terminal tyrosine residue is removed by vasohibin (VASH) carboxypeptidases [6,7] and MATCAP/TMCP family enzymes [8,9], and is re-ligated by a tubulin tyrosine ligase (TTL) [10]. This cycle plays key roles in neuronal transport, neurogenesis, myogenesis and cardiomyocyte contraction [11–14]. The tyrosinated and detyrosinated states of α-tubulin differentially regulate MAP interactions, motor protein behavior and MT properties. Tyrosination promotes MT dynamics and controls CAP-Gly protein binding [15], selective motor recruitment [16,17], kinetochore-MT attachment [18], dynein-dynactin motility [19], as well as retrograde intraflagellar transport (IFT) along the A-tubule [20,21]. Conversely, detyrosination accumulates on stable MTs, inhibits depolymerization by the kinesin-13 family [22], enhances cytoskeletal stiffness [12,23,24], and supports anterograde IFT along the B-tubule [20].

Strikingly, β-tubulin also carries the very C-terminal tyrosine in trypanosomatids, a feature of unknown biological function that is absent from other model eukaryotes [3]. This tyrosine is also subject to detyrosination [25], but unlike α-tubulin, β-tubulin does not undergo post-translational retyrosination in *T. brucei* [26]. Previous studies demonstrated that the VASH enzyme catalyses detyrosination of both α- and β-tubulin in *T. brucei* and *Leishmania* [27–29]. β-tubulin is nevertheless detyrosinated less efficiently than α-tubulin [29,30], indicating that the two subunits are differentially regulated. Importantly, the simultaneous alteration of α- and β-tubulin tyrosination status in previous functional studies has precluded a direct assessment of the specific role of β-tubulin. Consequently, it remains unknown whether β-tubulin tyrosination simply mirrors the well-established α-tubulin cycle or instead constitutes an independent regulatory layer of the tubulin code.

Another hallmark of trypanosomatid MTs is their extensive glutamylation, which involves the generation of glutamate side chains at the tubulin C-terminal tails catalyzed by multiple tubulin tyrosine ligase-like (TTLL) enzymes [25,30,31]. Several TTLL enzymes regulate cytoskeletal architecture and flagellar motility [32,33], and recent studies revealed a crosstalk between glutamylation and detyrosination [28–30,33]. Increased tubulin tyrosination is associated with reduced glutamylation, whereas enhanced detyrosination promotes glutamylation [28–30]. However, because functional studies simultaneously modified both tubulin subunits, whether this crosstalk is controlled by α-, β- or by both tubulin subunits remains unresolved.

Here, we combined comparative genomics, high-resolution imaging and precise genome editing to genetically uncouple α- and β-tubulin tyrosination in *T. brucei*. We show that the presence of a final tyrosine on β-tubulin is an evolutionarily conserved feature of trypanosomatids that is functionally distinct from the canonical α-tubulin tyrosination cycle. By selectively manipulating the tyrosination state of each tubulin subunit, we demonstrate that MT glutamylation is determined by the combined tyrosination status of α- and β-tubulin rather than by either subunit alone. Finally, we show that reciprocal β-tubulin tyrosination states reshape parasite morphology, single-cell motility and collective migration. Together, these findings establish β-tubulin tyrosination as a distinct regulatory layer in the trypanosomatid tubulin code that coordinates MT glutamylation and MT-dependent cellular functions.

## Results

### β-tubulin tyrosination is an evolutionarily conserved feature of trypanosomatids

To explore the evolutionary distribution of α- and β-tubulin tyrosination, we analyzed 504 eukaryotic genomes using the OrthoMCL [34] and EggNOG [35] databases, which include representative species from major eukaryotic supergroups: Opisthokonta, Archaeplastida, SAR (stramenopiles, alveolates, rhizaria), Amoebozoa and Excavata. We first generated a heatmap summarizing the presence or absence of C-terminal tyrosine motifs on α- and β-tubulin and mapped the occurrence of VASH and MATCAP/TMCP homologs (**Fig. S1**). These data were also summarized in a simplified, non-scaled phylogenetic tree to visualize the taxonomic distribution of the tubulin C-terminal motif (**Fig. 1A**).

**Figure 1.**
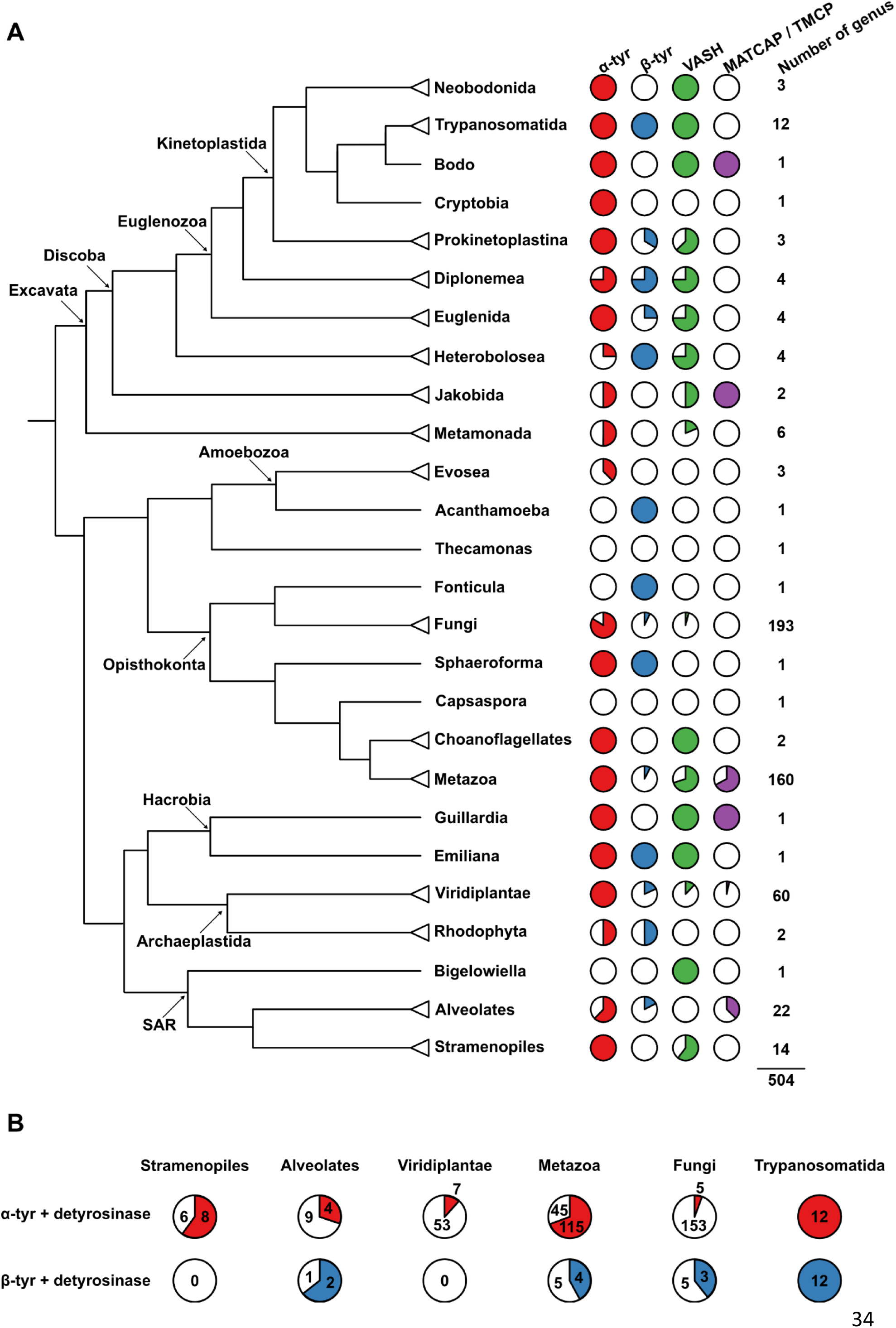
β-tubulin tyrosination is an evolutionarily conserved feature of trypanosomatids. (**A**) Phylogenetic representation summarizing, for each major eukaryotic clade, the presence of C-terminal tyrosine motifs in α- and β-tubulin and of the tubulin detyrosinases VASH and MATCAP/TMCP. The complete genus-level analysis and corresponding orthogroup assignments are presented in Fig. S1. (**B**) Co-occurrence analysis of an encoded C-terminal tyrosine motif and its corresponding detyrosinase, shown separately for α-tubulin and β-tubulin. Colors show positive co-occurrence indicating taxa in which both the tubulin tyrosine motif and the corresponding detyrosinase ortholog were detected, absence of colors indicate the presence of the tyrosine motif without corresponding enzyme.

The C-terminal tyrosine of α-tubulin was found to be widely conserved, with almost 90% of genera (n=445/504) containing this residue in at least one α-tubulin isotype. This high prevalence was consistent across all major clades, supporting α-tubulin tyrosination as an ancestral and conserved feature (**Fig. 1A and S1**). However, the presence of VASH and/or MATCAP/TMCP homologs was heterogeneous. Among Opisthokonta, more than 70% of metazoan genera (n=115/160) encoded a putative detyrosinase against only 3% of fungal genera (n=6/193). To determine whether the presence of a C-terminal tyrosine motif was associated with the predicted capacity for tubulin detyrosination, we assessed the co-occurrence of each motif with either VASH or MATCAP/TMCP homolog (**Fig. 1B**). To limit the influence of poorly represented lineages, this analysis was restricted to taxonomic groups represented by more than ten genera (Stramenopiles, Alveolates, Viridiplantae, Metazoa, Fungi and Trypanosomatida). Consistent with the broad conservation of the α-tubulin C-terminal tyrosine, its co-occurrence with a putative detyrosinase was frequent in most of these groups. This association was particularly prevalent in metazoans and Trypanosomatida, and remained detectable, although less frequent, in Alveolates and Stramenopiles (**Fig. 1B**). By contrast, the co-occurrence of a C-terminal tyrosine on α-tubulin and an identifiable VASH or MATCAP/TMCP homolog was rare in Viridiplantae and even less frequent in Fungi (**Fig. 1B**), indicating either the absence of detyrosination or the existence of alternative detyrosinases.

In contrast, the β-tubulin C-terminal tyrosine showed a markedly more restricted distribution and was detected in 11% of genera (n=56/504) overall (**Fig. 1A and S1**). The motif was absent from Stramenopiles and occurred in only a limited number of Alveolates, Opisthokonta, Amoebozoa and Viridiplantae genera. In these lineages, co-occurrence of β-tubulin tyrosination with a putative detyrosinase was variable: it was observed in some Alveolates, Metazoa and Fungi, but not in Viridiplantae (**Fig. 1B**). However, because β-tyr-positive genera were rare within these groups, these co-occurrence patterns were based on relatively small numbers of genera. Among excavates, on the other hand, β-tubulin tyrosination was consistently present in all Trypanosomatida included in the databases (*Angomonas; Blastocrithidia; Blechomonas; Crithidia; Endotrypanum; Leishmania; Leptomonas; Novymonas; Paratrypanosoma; Porcisia; Strigomonas; Trypanosoma*). Moreover, all β-Tyr-positive trypanosomatids only encoded a VASH ortholog (**Fig. 1A-B and S1**). Notably, the terminal tyrosine on β-tubulin was absent in *Bodo saltans,* the closest known non-parasitic relative of trypanosomatids [36]. Extending this analysis to a broader sampling of Euglenozoa revealed that the absence of a β-tubulin C-terminal tyrosine was in fact widespread across bodonids. Nevertheless, co-occurrence of the tyrosinated β-tubulin C-terminal tyrosine with a VASH homolog was detected in other euglenozoan lineages, including the prokinetoplastid *Apiculatamorpha* and the diplonemids *Diplonema*, *Hemistasia* and *Rhyncopus*. Outside Euglenozoa, this co-occurrence was also detected in the heteroloboseans *Pharyngomonas*, *Naegleria* and *Willaertia* (**Fig. S1**).

Together, these data revealed a clear evolutionary contrast between the broad conservation of α-tubulin tyrosination and the restricted distribution of the β-tubulin C-terminal tyrosine, whose co-occurrence with a putative detyrosinase was most prominent in trypanosomatids. Since β-tubulin detyrosination occurs in both *Leishmania* spp. and *T. brucei* [27,29] but only *T. brucei* expresses a single β-tubulin isotype [3,5], we selected this species as a tractable model to investigate the biological function of β-tubulin tyrosination in trypanosomatids.

### Distinct subcellular distribution of α- and β-tubulin tyrosination in *T. brucei*

The α-tubulin tyrosination cycle has long been characterized in *T. brucei* using the YL1/2 antibody, which recognizes tyrosinated α-tubulin [26,37,38]. To investigate the subcellular distribution of tyrosinated β-tubulin in comparison with α-tubulin, we generated a specific anti-Tbβtyr antibody for co-labeling analysis. We employed ultrastructure expansion microscopy (UExM) [39] to examine MT tyrosination profiles in procyclic trypanosomes using antibodies against tyrosinated α-tubulin (YL1/2) and β-tubulin (βtyr), as well as previously validated antibodies against detyrosinated α-tubulin (αΔ1) and β-tubulin (βΔ1) [27,28].

During interphase, βtyr staining largely overlapped with the YL1/2 pattern, with strong signals at the posterior cell body, distal flagellar portion, basal bodies and the MtQ (**Fig. 2A, upper panel, and 2B**). However, owing to its cross reactivity with TbRP2 [40], the YL1/2 signal extended more broadly around the mature basal body (**Fig. 2B**), consistent with previous immunogold labelling in this region [41]. Notably, βtyr labeling was stronger at the pro-basal body than at the mature basal body (**Fig. 2B**), indicative of active detyrosination during basal body maturation.

**Figure 2.**
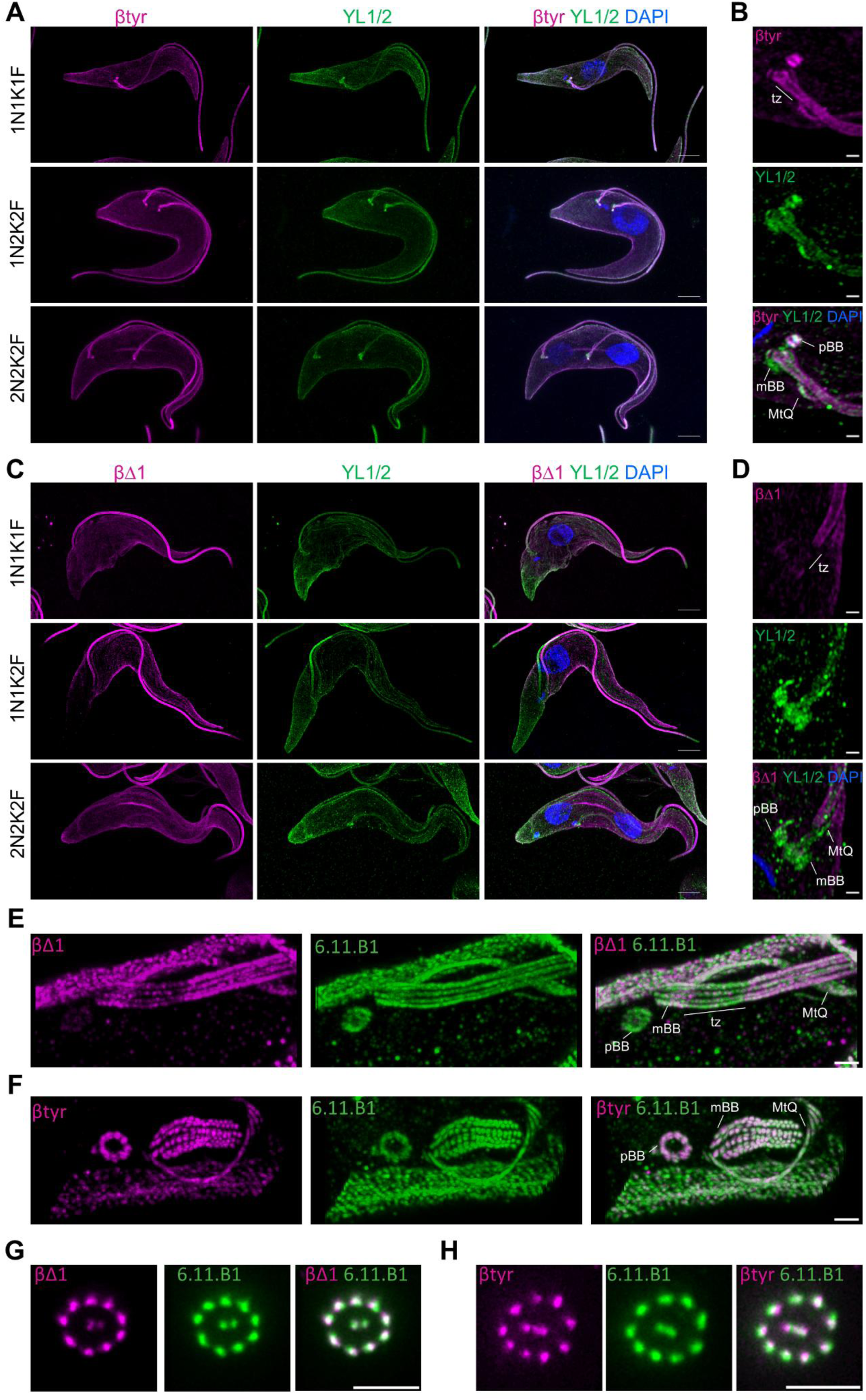
Distinct subcellular distribution of α- and β-tubulin tyrosination in *T. brucei*. (**A-D**) UexM analysis of tyrosinated and detyrosinated α- and β-tubulin in *T. brucei* parental procyclics. Maximum intensity projections of z-stack confocal images are presented. The αΔ1 and YL1/2 antibodies recognize detyrosinated or tyrosinated α-tubulin, respectively. The βΔ1 and β-tyr antibodies recognize detyrosinated or tyrosinated β-tubulin, respectively. White arrows indicate the mitotic spindle. p : posterior; a : anterior; of: old flagellum; nf: new flagellum; tz: transition zone; mBB: mature basal body; pBB: pro-basal body; MtQ: microtubule quartet. Scale bars: 10 µm (A and C) or 1µm (B and D). (**E-F**) iUexM analysis of detyrosinated (E) and tyrosinated (F) β-tubulin in *T. brucei* parental procyclics. Maximum intensity projections of z-stack confocal images are presented. Scale bar: 2 µm. (**G-H**) iUexM analysis of detyrosinated (G) and tyrosinated (H) β-tubulin in *T. brucei* parental procyclics in axonemal cross sections. Single Z-stack are presented. Scale bar: 2 µm.

Consistent with the distribution of tyrosinated tubulin, detyrosinated α-tubulin (αΔ1) displayed a complementary pattern, being absent from the posterior cell body and enriched along the flagellum (**Fig. S2A, upper panel**). While the transition zone above the mature basal body lacked αΔ1 signal, both basal bodies were labelled, with a slightly stronger staining at the mature basal body (**Fig. S2B**), in line with the higher level of tyrosinated tubulin at the pro-basal body (**Fig. 2B**). Detyrosinated β-tubulin (βΔ1) exhibited a distribution similar to αΔ1 throughout the cell body and flagellum (**Fig. 2C**), with no signal in the transition zone and only weak labeling at basal bodies (**Fig. 2D**).

To further resolve the distribution of tyrosinated and detyrosinated β-tubulin from the basal body to the axoneme, we combined iterative U-ExM with STED microscopy, using acetylated α-tubulin (6.11.B1) as a structural reference (**Fig. 2E-H**). βΔ1 labeling at the mature basal body ranged from weak to clearly detectable, although the most proximal region remained unlabeled (**Fig. 2E and S2C**). However, it was barely present at the pro-basal body or within the transition zone (**Fig. 2E**). By contrast, βtyr strongly labeled both basal bodies, with marked enrichment at the pro-basal body (**Fig. S2D-E**), and extended from the mature basal body triplets into the transition zone (**Fig. 2F**). In transverse axonemal sections, both βtyr and βΔ1 labeled all nine outer doublets (**Fig. 2G-H**), whereas the central pair was more prominently labeled by βtyr (**Fig. 2G**). Within individual outer doublets, βtyr and βΔ1 labeling appeared preferentially associated with one of the two tubules (**Fig. 2G-H**), revealing differential β-tubulin tyrosination between the A- and B-tubules. These observations indicate that the central pair remains largely tyrosinated and reveal β-tubulin tyrosination heterogeneity within the axoneme.

During progression through the cell cycle, the redistribution of tyrosinated and detyrosinated tubulins revealed further differences between α- and β-tubulin. In both subunits, tyrosination was enriched at sites of active MT assembly, such as the growing posterior corset and the emerging flagellum (**Fig. 2A, middle panel**), whereas the corresponding detyrosinated forms were abundant along mature MT arrays (**Fig. 2C and S2A, middle panels**). However, β-tubulin showed a broader, more persistent tyrosination pattern than α-tubulin, with βtyr extending throughout the cell body and along the full length of both mature and growing flagella (**Fig. 2A, middle panel**). In addition, while αΔ1 (**Fig. S2A, lower panel**) and βΔ1 (**Fig. 2C, lower panel**) both labelled the mitotic spindle, βtyr, unlike YL1/2, also marked spindle MTs (**Fig. 2A, lower panel).** This observation is consistent with recent proteomic analyses [30] and with the slower VASH-mediated detyrosination kinetics reported for β-tubulin in *Leishmania* [29].

Altogether, these observations reveal distinct subcellular distributions of α- and β-tubulin tyrosination, with β-tubulin maintaining a broader tyrosinated state throughout the cell cycle, consistent with differential detyrosination of the two subunits.

### Engineering a constitutively detyrosinated β-tubulin state

In *T. brucei*, α- and β-tubulin are encoded as alternating tandem repeats on chromosome 1 [5]. To investigate the functional contribution of β-tubulin tyrosination-detyrosination cycle, we employed a recently developed CRISPR-Cas9 ribonucleoprotein system [42] to simultaneously edit all β-tubulin gene copies and introduce a donor sequence lacking the terminal tyrosine codon (**Fig. S3A**). Therefore, we genetically engineered a constitutively detyrosinated β-tubulin state while leaving the α-tubulin intact (**Fig. 3A**). Editing was attempted in both monomorphic Tb427 and pleomorphic, differentiation-competent, Tb927 cell lines. However, complete deletion of the C-terminal tyrosine was only obtained in a single Tb427 clone, which was confirmed by Sanger and Nanopore sequencing (**Fig. S3B**). As such, the Tb427 clone was used for all subsequent analyses (hereafter referred to as βΔ1). Nanopore sequencing (**Fig. S3C**) and qPCR (**Fig. S3D**) revealed a concomitant reduction in tubulin gene copy number following homologous repair. Nonetheless, immunoblot analysis showed similar tubulin protein levels relative to the parental cell line (**Fig. S3E**), indicating post-transcriptional compensation of tubulin abundance in the βΔ1 mutant.

**Figure 3.**
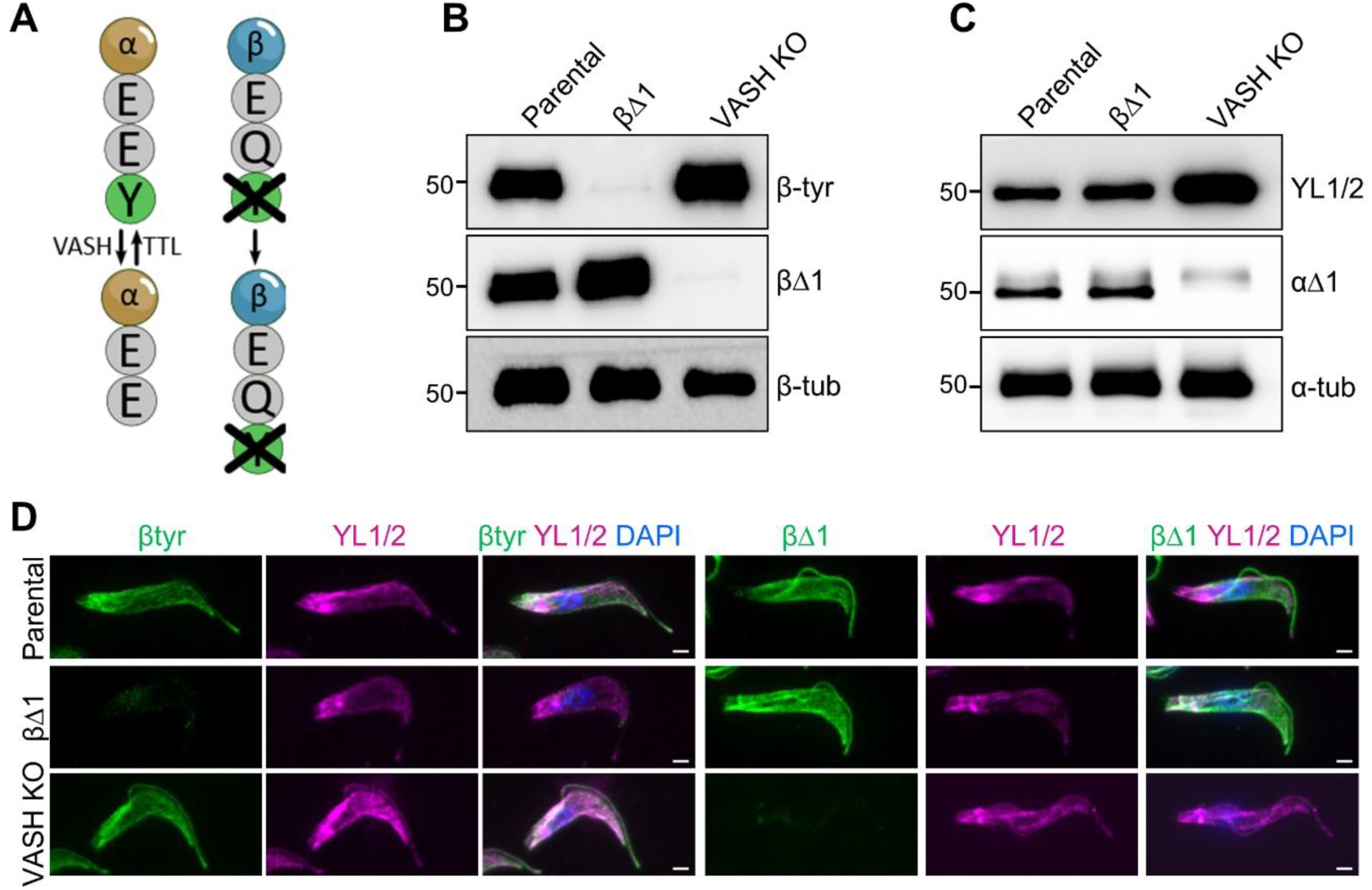
Engineering a constitutively detyrosinated β-tubulin state. (**A**) Schematic representation of the tyrosination status of the βΔ1 mutant cell line. (**B**) Immunoblot analysis of β-tubulin tyrosination levels in whole cell extracts from parental, βΔ1 mutant and VASH KO cell lines. β-tubulin is used as a loading control. (**C**) Immunoblot analysis of α-tubulin tyrosination levels in whole cell extracts from parental, βΔ1 mutant and VASH KO cell lines. α-tubulin is used as a loading control. (**D**) Immunofluorescence analysis of tyrosinated and detyrosinated α- and β-tubulin of detergent-extracted cytoskeletons from parental, βΔ1 mutant and VASH KO cell lines. Scale bar: 2µm.

As a complementary condition in which tubulins remain constitutively tyrosinated [27,28], we generated a VASH knockout (VASH KO) cell line in the same Tb427 background using CRISPR-Cas9 (**Fig. S3F**). Immunoblot analysis confirmed the absence of tyrosinated β-tubulin in the βΔ1 mutant and a corresponding increase in the detyrosinated form (**Fig. 3B**). In contrast, VASH KO cells displayed the opposite profile, with fully tyrosinated α- and β-tubulin (**Fig. 3B and 3C**), as previously reported [27,28]. Importantly, α-tubulin tyrosination remained unchanged in the βΔ1 mutant as compared with the parental cell line (**Fig. 3C**), indicating that α-tubulin de-/re-tyrosination cycles proceed normally upon constitutive β-tubulin detyrosination. Consistently, immunofluorescence analysis of detergent-extracted cells showed the presence of detyrosinated β-tubulin throughout the cell body accompanied by the loss of the tyrosinated form, whereas α-tubulin tyrosination remained restricted to the posterior cell body and distal flagellar tip, similar to the parental line (**Fig. 3D**).

Next, we assessed βΔ1 mutant proliferation over four days. Early-passage βΔ1 cells displayed a growth defect following clone isolation, whereas cells maintained in culture for approximately two months proliferated at rates comparable to those of the parental line (**Fig. S3G**). To test whether this transient growth defect was associated with cell cycle perturbations, we stained the nucleus (N) and the kinetoplast (K) (mitochondrial DNA) with Hoechst, two organelles whose division is tightly coordinated during parasite replication [43]. The early-passage βΔ1 population displayed a slight reduction in interphasic (1N1K) cells and an increase in aberrant configurations, including zoids (0N1K) and binucleated cells (2N1K) (**Fig. S3H**). These abnormalities were no longer present in the long-term cultured population, indicating a compensatory response over time. The levels of tubulin remained unchanged between early and long-term cultured βΔ1 cells (**Fig. S3E**), excluding compensation through tubulin overexpression. Likewise, β-tubulin remained constitutively detyrosinated (**Fig. S3I**), indicating that the recovery of normal proliferation did not result from β-tubulin retyrosination, consistent with previous reports showing that this subunit, in contrast to α-tubulin, is not a substrate of TTL [26].

Altogether, these results demonstrate that constitutive β-tubulin detyrosination does not perturb the α-tubulin tyrosination cycle but transiently compromises cell proliferation and cell cycle progression. These defects are subsequently rescued by compensatory mechanisms that occur independently of β-tubulin retyrosination or tubulin overexpression.

### Engineering selective inhibition of β-tubulin detyrosination

To specifically test the function of β-tubulin detyrosination, we sought to inhibit VASH-mediated detyrosination of this subunit. A single amino acid substitution in human α-tubulin, replacing the penultimate glutamate residue with an aspartate (GEEY → GEDY), was previously shown to inhibit VASH activity [8]. We therefore introduced the equivalent mutation in β-tubulin, substituting the penultimate glutamine with aspartate (EEQY → EEDY) to selectively inhibit β-tubulin detyrosination without altering the α-tubulin tyrosination status (**Fig. 4A**). Using the same marker-free CRISPR-Cas9 strategy as for the βΔ1 mutant (**Fig. S3A**), we generated a clonal βQ441D mutant in which all β-tubulin gene copies carried the intended substitution, as confirmed by Sanger sequencing (**Fig. S4A**). qPCR showed no major change in tubulin gene copy number (**Fig. S4B**) or total protein levels (**Fig. S4C**).

**Figure 4.**
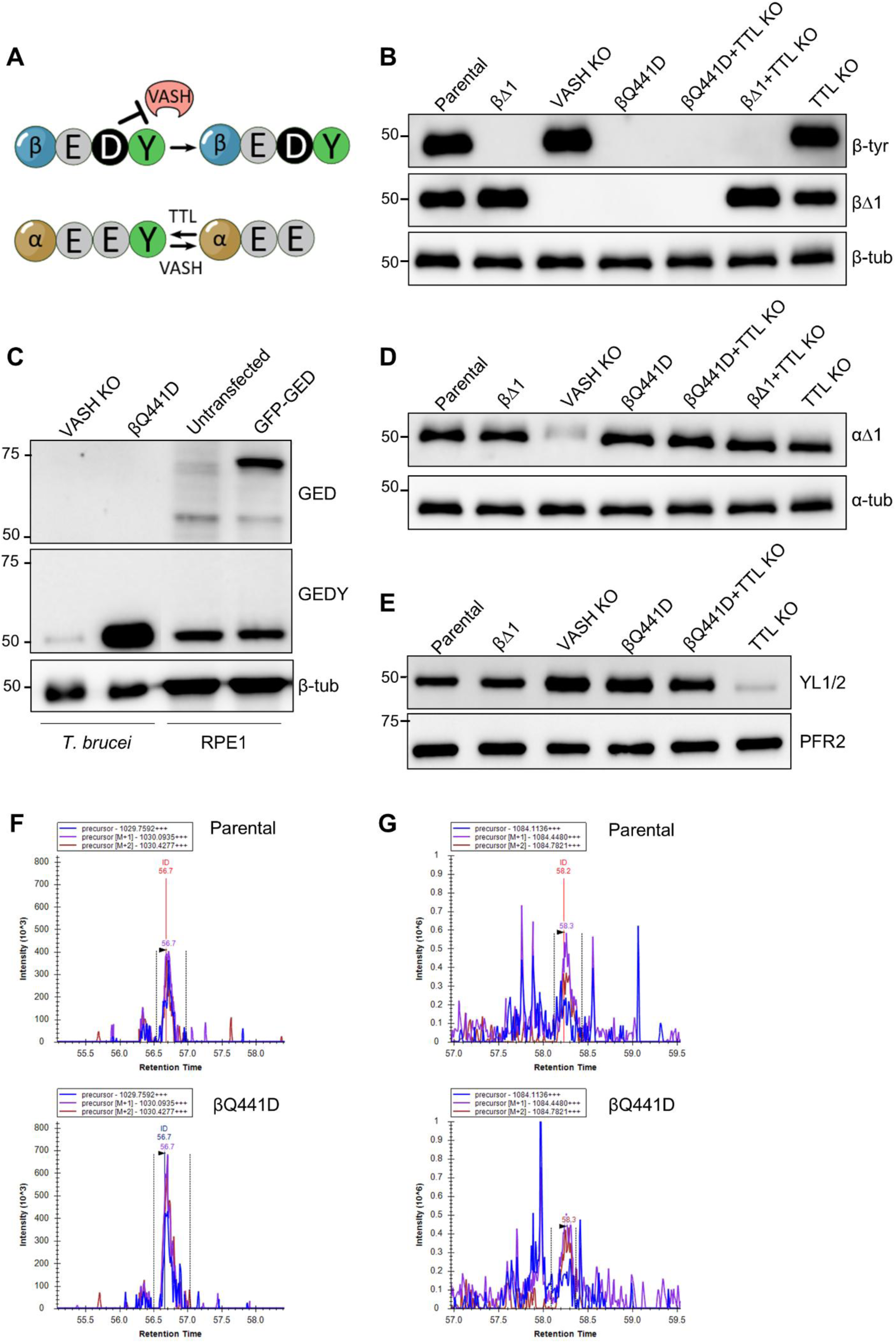
Engineering selective inhibition of β-tubulin detyrosination. (**A**) Schematic representation of the tyrosination status of the βQ441D mutant cell line in which VASH-mediated detyrosination is impaired. (**B**) Immunoblot analysis of β-tubulin tyrosination levels in whole cell extracts from parental, βΔ1 mutant, VASH KO, βQ441D mutant, TTL KO, βQ441D TTL KO and βΔ1 TTL KO cell lines. β-tubulin is used as a loading control. (**C**) Immunoblot analysis of VASH-mediated β-tubulin detyrosination in whole-cell extracts from VASH KO, and βQ441D mutant cell lines using GEDY and GED antibodies. Whole-cell extracts from control (untransfected) and GFP-GED-expressing RPE1 cells were included as negative and positive control for antibody specificity, respectively. GFP-GED (∼75 kDa) is recognized by GED but not by GEDY. β-tubulin is used as a loading control. (**D**) Immunoblot analysis of detyrosinated α-tubulin levels in whole cell extracts from parental, βΔ1 mutant, VASH KO, βQ441D mutant, TTL KO, βQ441D TTL KO and βΔ1 TTL KO cell lines. α-tubulin is used as a loading control. (**E**) Immunoblot analysis of tyrosinated α-tubulin levels in whole cell extracts from parental, βΔ1 mutant, VASH KO, βQ441D mutant, TTL KO, βQ441D TTL KO and βΔ1 TTL KO cell lines. PFR2 is used as a loading control. (**F-G**) Mass spectrometry analysis of α-tubulin C-terminal peptides from parental and βQ441D mutant cell lines. Representative extracted ion chromatograms corresponding to detyrosinated (F) and tyrosinated (G) α-tubulin are shown.

Immunoblot analyses showed that neither βΔ1 nor βtyr signal was detectable in βQ441D cells (**Fig. 4B**), suggesting that Q → D substitution disrupts the epitopes recognized by these antibodies. We therefore used antibodies against the GEDY and GED motifs [8] to directly determine whether the βQ441D substitution prevents β-tubulin detyrosination. βQ441D cells displayed high levels of tyrosinated β-tubulin, whereas no detyrosinated β-tubulin was detected (**Fig. 4C**). The absence of a signal for the anti-GED antibody, despite the recognition of the EED motif in GFP-EED-expressing RPE1 cells (**Fig. 4C**), confirmed that the βQ441D substitution prevents VASH-mediated detyrosination of β-tubulin. In this background, detyrosinated α-tubulin levels remained unchanged (**Fig. 4D**), whereas apparent levels of tyrosinated α-tubulin increased (**Fig. 4E),** suggesting either enhanced YL1/2 affinity for the EDY motif or altered α-tubulin tyrosination.

To distinguish between altered epitope recognition and genuine changes in α-tubulin tyrosination, we deleted TTL, the α-tubulin retyrosination enzyme, in both parental and βQ441D backgrounds (**Fig. S4D**). As expected, β-tubulin signals remained unchanged in the TTL KO cells (**Fig. 4B**), confirming that TTL does not modify β-tubulin [26]. Similarly, βQ441D TTL KO cells displayed the same β-tubulin profile as βQ441D cells (**Fig. 4B**). In contrast, tyrosinated α-tubulin was nearly completely abolished in TTL KO cells and only partially reduced in the double mutant (**Fig. 4D**). Further resolving the YL1/2 signal revealed an additional band in both the single βQ441D and the double βQ441D TTL KO mutants (**Fig. S4E**). Together, these observations support the idea that the elevated YL1/2 signal observed in βQ441D cells reflects EDY epitope recognition rather than increased α-tubulin tyrosination. To independently validate this conclusion, we performed targeted LC-MS/MS analyses of tubulin C-terminal peptides. Extracted ion chromatograms of detyrosinated (**Fig. 4E**) and tyrosinated (**Fig. 4F**) α-tubulin C-terminal peptides, as well as internal α-tubulin peptides (**Fig. S4F**), showed comparable signal intensities in parental and βQ441D samples, confirming that the βQ441D substitution does not alter the α-tubulin tyrosination cycle.

Next, we assessed proliferation of the different cell lines over four days. βQ441D and VASH KO cells proliferated at rates comparable to the parental control (**Fig. S4G**), consistent with previous observations in monomorphic *T. brucei* [28], as well as in *Leishmania mexicana* promastigotes [29]. Similarly, TTL KO and βQ441D TTL KO double mutant cells displayed normal proliferation (**Fig. S4G**).

Taken together, these results establish βQ441D as a genetic tool that selectively blocks β-tubulin detyrosination without perturbing the α-tubulin tyrosination cycle, demonstrating that the two tubulin subunits can be independently regulated despite sharing the same enzymatic machinery.

### β-tubulin tyrosination independently regulates microtubule glutamylation

Recent work in *T. brucei* showed that detyrosination and polyglutamylation are coupled through a positive feedback loop, whereby detyrosination promotes polyglutamylation, which in turn enhances further detyrosination [28,30]. To determine whether β-tubulin tyrosination influences glutamylation, we analyzed a panel of cell lines displaying distinct α/β tyrosination states: parental cells (α- and β-tubulin tyrosinated/detyrosinated), βΔ1 (β-detyrosinated), TTL KO (α-detyrosinated), βQ441D (β-tyrosinated), βQ441D TTL KO (α-detyrosinated/β-tyrosinated), and VASH KO (α- and β-tyrosinated).

We performed immunoblot analyses using GT335 (branching-point glutamate), PolyE (long glutamate chains), and βmonoE (βE435 branch point) [44] antibodies. VASH KO and βQ441D cells, lacking detyrosinated β-tubulin (**Fig. 4B**), displayed reduced GT335 and PolyE signals compared to parental cells (**Fig. 5A and 5B**), as previously shown in *Leishmania mexicana* [29] and *T. brucei* [28]. The reduction was strongest in VASH KO cells, in which both α- and β-tubulin remain tyrosinated, whereas βQ441D cells exhibited an intermediate decrease, indicating that maintenance of β-tubulin tyrosination alone partially reduces glutamylation.

**Figure 5.**
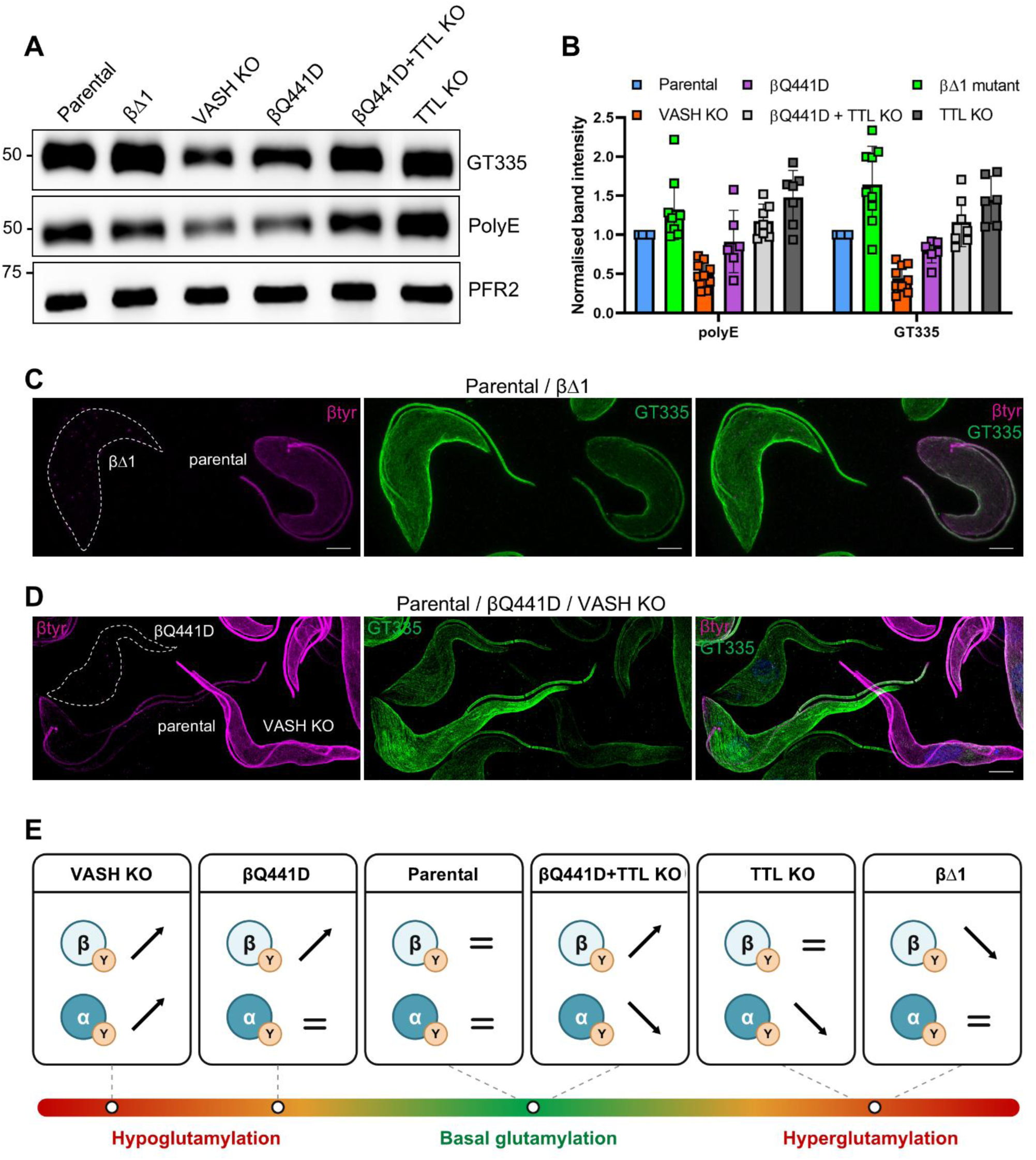
β-tubulin tyrosination independently regulates microtubule glutamylation. (**A**) Immunoblot analysis of microtubule glutamylation in whole-cell extracts from parental, βΔ1 mutant, VASH KO, βQ441D mutant, TTL KO and βQ441D TTL KO cell lines. The GT335 antibody recognizes glutamate side chains of any length. The PolyE antibody recognizes glutamate side chains of at least four glutamate residues. PFR2 is used as a loading control. (**B**) Quantification of immunoblot signals detected with GT335 and PolyE antibodies. Band intensities were normalized to the loading control and expressed relative to the parental cell line. Bars represent the mean ± SD from six independent experiments. (**C-D**) UexM analysis of microtubule glutamylation of parental and βΔ1 mutant cell lines or parental, VASH KO and βQ441D embedded in the same gel. Maximum intensity projections of z-stack confocal images are presented. Scale bar : 10µm. (**E**) Proposed model summarizing the levels of tubulin glutamylation observed in the indicated cell lines showing that modulation of either α- or β-tubulin tyrosination independently affects the level of tubulin glutamylation.

Conversely, βΔ1 cells, lacking tyrosinated β-tubulin (**Fig. 3B and 4B**), displayed a pronounced increase in GT335 signal and a more moderate increase in PolyE signal as compared to parental cells (**Fig. 5A and B**). Although less marked, the increase in PolyE was reproducibly detected and confirmed using serial dilutions of protein extracts (**Fig. S5A**). To determine whether this increase was associated with the initial growth defect of the βΔ1 mutant (**Fig. S3G**), we compared early-passage and long-term cultured βΔ1 cell populations. Both showed increased GT335 and PolyE signals relative to parental cells (**Fig. S5B**), indicating that enhanced glutamylation was associated with constitutive β-tubulin detyrosination rather than with the growth defect itself. Similarly, TTL KO cells, characterized by extensive α-tubulin detyrosination, also displayed elevated glutamylation levels (**Fig. 5A and 5B**). To determine whether simultaneous detyrosination of both α- and β-tubulin further enhances glutamylation, we analyzed the βΔ1 TTL KO double mutant which did not show any detectable increase in GT335 and PolyE signals as compared to the single βΔ1 mutant (**Fig. S5C**). The βQ441D TTL KO double mutant, which combines constitutively tyrosinated β-tubulin with detyrosinated α-tubulin, exhibited glutamylation levels comparable to the parental line, indicating that opposing α- and β-tubulin tyrosination states restore intermediate glutamylation levels (**Fig. 5A and 5B**). Importantly, βmonoE signal intensity remained unchanged in VASH KO and βΔ1 cells (**Fig. S5A**), indicating that the changes observed with GT335 and PolyE reflect altered glutamylation at other sites and/or glutamate chain length rather than differences in βE435 occupancy.

We next performed expansion microscopy on mixed cell populations, distinguishing cell lines using the β-tyr antibody. Four combinations were analyzed: (1) parental and βΔ1 (**Fig. 5C**); (2) parental, βQ441D and VASH KO (**Fig. 5D**); (3) βΔ1 and TTL KO (**Fig. S5D**); and (4) parental and βQ441D TTL KO double mutant (**Fig. S5E**).

Consistent with the immunoblot analyses (**Fig. 5A and 5B**), βΔ1 cells, characterized by a negative β-tyr signal, exhibited an overall increase in glutamylation intensity compared to parental cells (**Fig. 5C**). In images acquired separately for each cell line, GT335 signal was mostly excluded from basal bodies in parental cells, whereas it became clearly detectable at these structures in βΔ1 cells (**Fig. S5F**), indicating enhanced initiation of glutamylation upon constitutive β-tubulin detyrosination. For PolyE, both parental and βΔ1 cells displayed weak signal on posterior cortical MTs, enrichment at the pro-basal body and absence from the transition zone (**Fig. S5G**).

Analysis of the mixed population including parental, βQ441D and VASH KO cells confirmed the immunoblot results (**Fig. 5D**). GT335 labeling was reduced in βQ441D cells and was lowest in VASH KO cells as compared to parental cells, with the three cell lines readily distinguishable by βtyr staining. Likewise, βQ441D TTL KO cells showed glutamylation levels comparable to the parental line (**Fig. S5E**), whereas βΔ1 and TTL KO cells displayed similarly elevated GT335 labeling (**Fig. S5D**), supporting the conclusion that constitutive detyrosination of either α- or β-tubulin enhances glutamylation.

Taken together, these results support a model in which the level of tubulin glutamylation reflects the tyrosination status of both α- and β-tubulin rather than either subunit alone (**Fig. 5E**). Increased detyrosination of either β-tubulin (βΔ1) or α-tubulin (TTL KO) enhances glutamylation, whereas constitutive β-tubulin tyrosination (βQ441D) partially reduces this modification. When both α- and β-tubulin remain fully tyrosinated, as in VASH KO cells, glutamylation is most strongly reduced. Conversely, re-establishing a mixed tyrosination state in the βQ441D TTL KO double mutant restores glutamylation to near-parental levels. These findings indicate that α- and β-tubulin tyrosination each independently contribute to the overall regulation of MT glutamylation.

### β-tubulin tyrosination impacts parasite morphology

Previous studies reported that altering tubulin tyrosination can affect *T. brucei* morphology, with VASH deletion causing a marked cell-body elongation in pleomorphic procyclic forms [27], while only modest reductions in cell and flagellum length were observed in a monomorphic cell line [28], as well as in *Leishmania* stationary promastigotes [29]. We therefore asked whether selective manipulation of β-tubulin tyrosination affects parasite morphology in our panel of engineered mutants.

Scanning electron microscopy revealed differences in parasite morphology, particularly in the free distal portion of the flagellum, which appeared shorter in βΔ1 cells and longer in βQ441D cells compared with parental cells (**Fig. 6A**). To further characterize these differences, we measured several morphological parameters in exponentially growing and stationary-phase cultures. Cell body length increased in stationary-phase cultures across all cell lines, with no differences observed during exponential growth and only βQ441D cells displaying a shorter cell body than parental cells in stationary phase (**Fig. S6A**). Total flagellum length was comparable between parental, βΔ1, βQ441D and VASH KO cells within each growth phase, and also increased in stationary-phase cultures across all cell lines (**Fig. 6B and S6B**). Consistent with the scanning electron microscopy observations, measurements of the free distal flagellum revealed a shorter free portion in βΔ1 cells and a longer one in βQ441D cells compared with parental cells, in both growth phases (**Fig. 6C**). In VASH KO cells, the free flagellum was slightly shorter during exponential growth (**Fig. S6C**) but comparable to parental cells in stationary phase (**Fig. 6A and S6C**).

**Figure 6.**
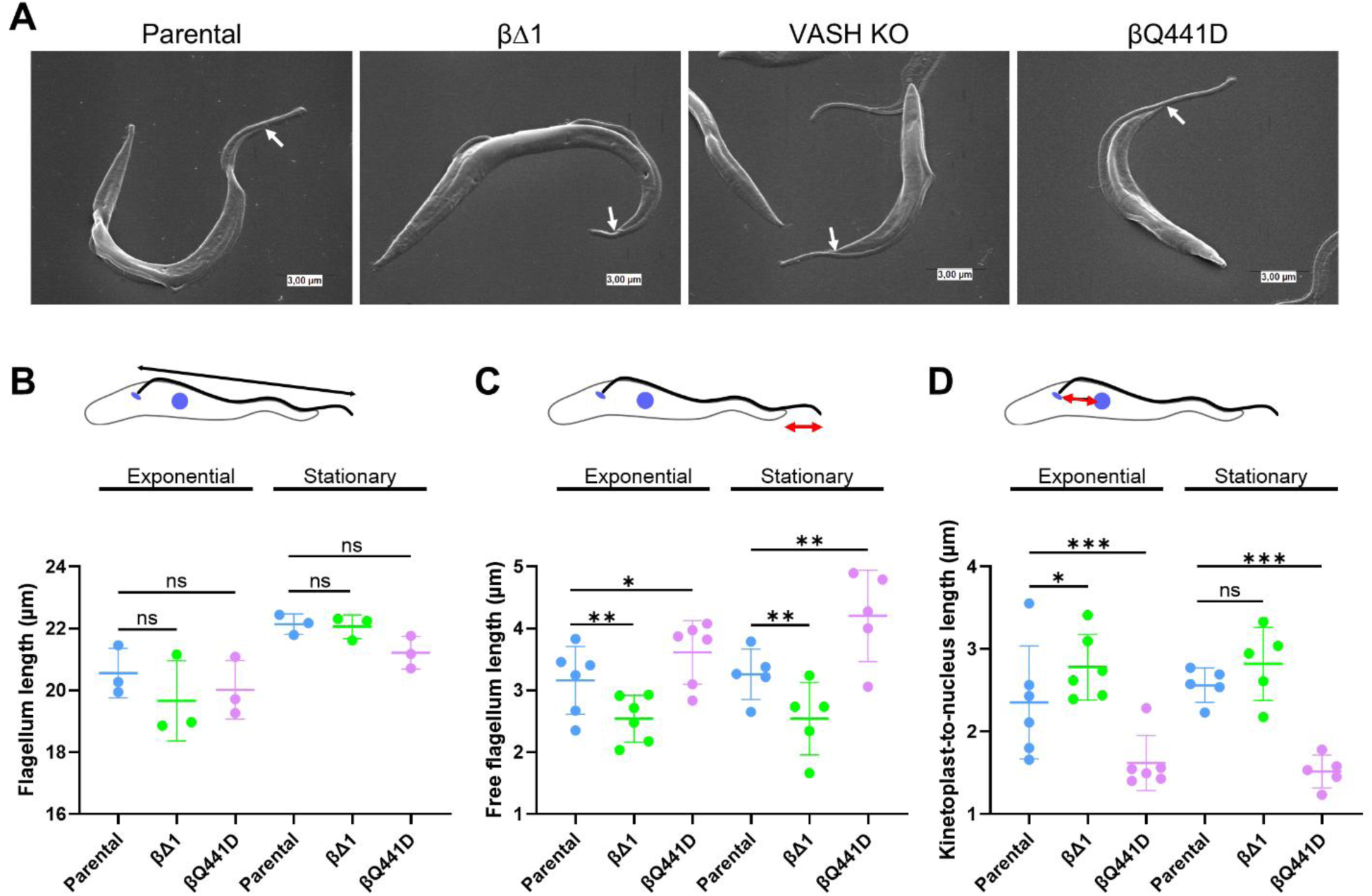
β-tubulin tyrosination impacts parasite morphology. (**A**) Scanning electron microscopy images of parental, βΔ1 mutant, VASH KO and βQ441D mutant cells in stationary phase, illustrating the shorter and longer free flagellum of the βΔ1 and βQ441D mutant cells, respectively. White arrows indicate where the free distal portion of the flagellum begins. Scale bar: 3 µm. (**B-D**) Measurements of total flagellum length (B), free flagellum length (C) and kinetoplast-to-nucleus distance (D) from parental, βΔ1 mutant and βQ441D mutant cell lines cultured in exponential and stationary phase. Each point represents the mean of one independent experiment (n=50 cells per experiment) and bars show the mean ± SD. Flagellum length was measured on mAb25-labelled cells (three independent experiments per growth phase). Free flagellum length and kinetoplast-to-nucleus distance were measured on mAb25- or PFR2-labelled cells (six independent experiments in exponential phase and five in stationary phase). Cells were labelled with β-tubulin for the cell body and Hoechst 33342. Each mutant cell line was compared with the parental cell line within the same growth phase using a repeated-measures one-way ANOVA followed by Dunnett’s multiple comparisons test (ns, not significant; *P<0.05; **P<0.01; ***P<0.001). The schematic above each graph indicates the parameter measured (highlighted by the arrow). Red arrows indicate parameters showing opposite changes in βΔ1 and βQ441D mutant cells.

To determine whether these changes resulted from altered flagellum assembly, we examined the growth of the new flagellum. In the grow-and-lock model, the flagellum elongates until a locking event, proposed to occur after cell division, prevents further elongation or shortening [45]. FLAM8 is abundant at the distal tip of mature flagella and progressively accumulates at the tip of the new flagellum as it elongates [45]. We therefore measured the ratio between the new and the old flagellum (NF/OF) in exponentially growing 2K2N cells with FLAM8 detected at the tips of both flagella. In parental cells, the NF/OF ratio was close to the expected ∼80% [45]. This ratio remained unchanged in all mutant backgrounds (**Fig. S6D**), indicating that the tyrosination status of β-tubulin does not affect new flagellum elongation.

We next measured the distance between the kinetoplast and the nucleus (**Fig. 6D and S6E**). During exponential growth, the kinetoplast-to-nucleus distance was increased in βΔ1 cells and reduced in βQ441D cells compared with parental cells (**Fig. 6D**). Similar trends were observed in stationary phase, although the difference reached statistical significance only in βQ441D cells (**Fig. 6D**). In VASH KO cells, this distance was also reduced during exponential growth, but did not differ significantly from that of parental cells in stationary phase (**Fig. S6E**). In contrast, the kinetoplast-to-posterior distance showed no significant differences between cell lines in either growth phase (**Fig. S6F**). Thus, βΔ1 and βQ441D cells displayed opposite changes in both free flagellum length and kinetoplast-to-nucleus distance.

Together, these results show that the tyrosination status of β-tubulin influences parasite morphology without altering new flagellum assembly. Constitutive β-tubulin detyrosination in βΔ1 cells was associated with a shorter free distal flagellum and an increased kinetoplast-to-nucleus distance, whereas βQ441D cells displayed the opposite changes in both growth phases.

### β-tubulin tyrosination controls parasite motility

Because trypanosome motility originates at the distal flagellar tip and propagates toward the cell body [46,47], we examined whether the morphological perturbations in the mutant cell lines altered swimming behavior. Trypanosome trajectories consist of alternating runs and tumbling events. Runs are characterized by persistent directional swimming, whereas tumbling events produce rapid changes in cell orientation before directional swimming resumes.

VASH KO cells swam at speeds comparable to the parental cell line, with no significant changes in run duration, directionality or tumbling frequency (**Fig. 7A-C**). In contrast, βΔ1 cells displayed a significant reduction in average swimming speed (**Fig. 7A**) and increased tumbling (**Fig. 7C**). βQ441D cells exhibited the opposite phenotype, with increased swimming speed (**Fig. 7A**) and reduced tumbling (**Fig. 7C**). While overall directionality was largely similar between mutants and parental cells, βQ441D cells showed significantly greater directional persistence as compared to βΔ1 cells (**Fig. 7B**).

**Figure 7.**
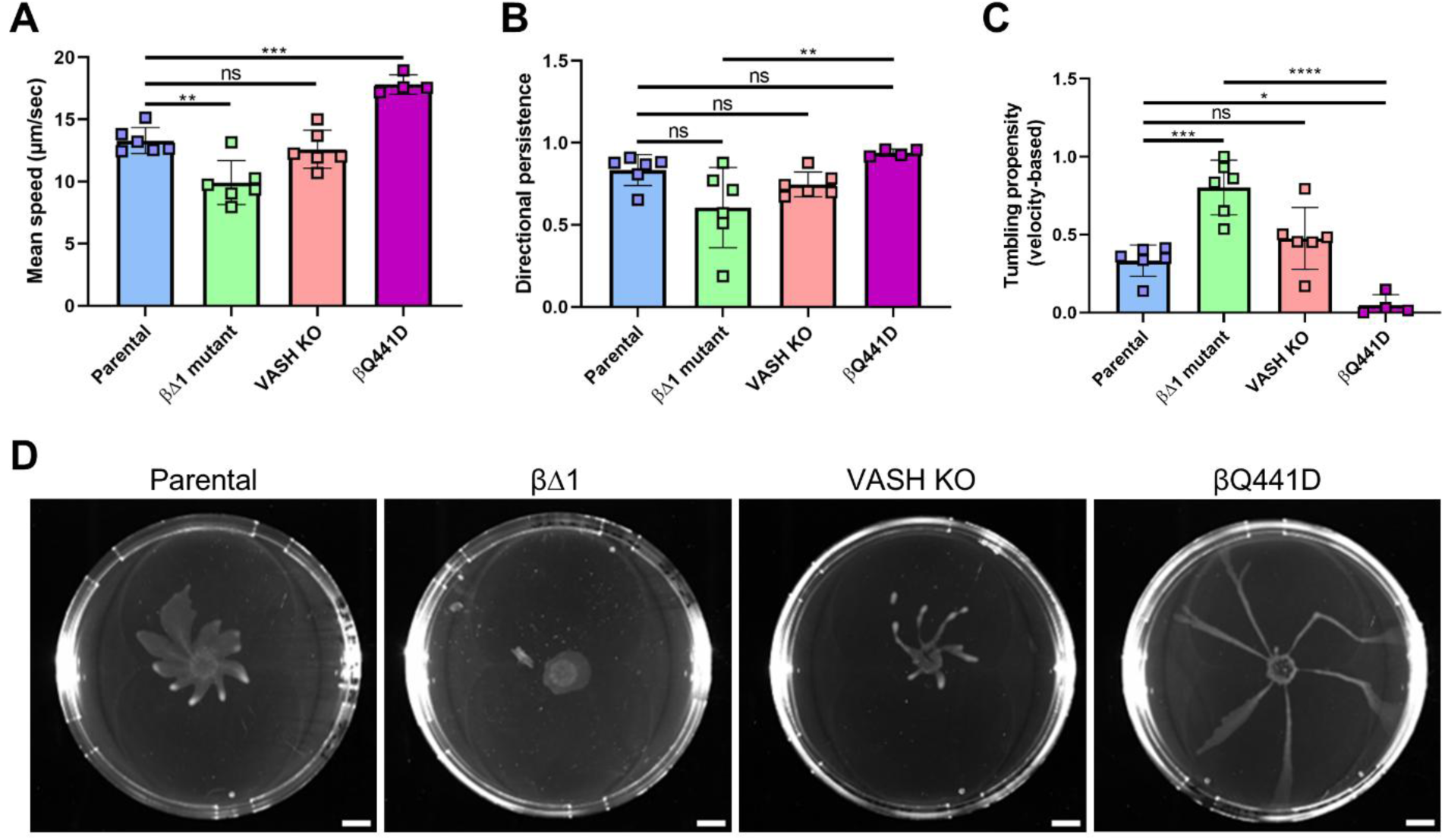
β-tubulin tyrosination controls parasite motility. (**A-C**) Swimming speed, directional persistence and tumbling propensity measurements from parental, βΔ1 mutant, VASH KO and βQ441D mutant cell lines obtained during exponential growth. (n= 4 from four independent experiments). (**D**) Photographs of social motility communities 4 days post-plating from parental, βΔ1 mutant, VASH KO and βQ441D mutant cell lines. Scale bar: 1cm.

Next, we investigated whether these changes in individual swimming behavior affected collective migration. Procyclic trypanosomes form organized migrating communities *in vitro*, which has been proposed to model colonization of the tsetse midgut [48]. We therefore evaluated social motility (SoMo) on semi-solid agar. Parental cells generated characteristic arrays of radial projections (**Fig. 7D**). Although thinner, projections from VASH KO cells were comparable in number and organization. In contrast, βΔ1 cells failed to generate projections and expanded isotropically as a homogeneous circular front (**Fig. 7D**), consistent with their reduced swimming speed and increased tumbling (**Fig. 7A and 7C**). Conversely, βQ441D cells formed straighter, more radially oriented projections with reduced curvature (**Fig. 7D**), consistent with their increased directional persistence at the single-cell level (**Fig. 7B**).

Altogether, modulation of the β-tubulin tyrosination-detyrosination balance produced opposite effects on single-cell and collective motility in *T. brucei*. While constitutive detyrosination slowed and reduced directional movement, inhibition of detyrosination increased swimming speed, trajectory persistence and collective organization. These findings indicate that β-tubulin tyrosination functions as a regulator of MT-dependent motile behaviours.

## Discussion

Tubulin detyrosination is a highly conserved component of the tubulin code, yet it has been almost exclusively considered in the context of the canonical α-tubulin tyrosination cycle. Trypanosomatids represent a particularly informative model to revisit this PTM. First, their MT-rich cytoskeleton relies on a limited repertoire of α- and β-tubulin isotypes, making PTMs a major source of MT diversity [3]. More strikingly, these parasites express β-tubulin with a C-terminal tyrosine which is subjected to detyrosination [25]. The functional significance of this modification, however, has remained unknown. Here, by selectively manipulating the β-tubulin tyrosination state in *T. brucei*, we show that it constitutes an additional level of regulation within the trypanosomatid tubulin code which contributes to the control of MT glutamylation, parasite morphology and motility in procyclics.

The function of β-tubulin tyrosination should be considered in the context of its heterogeneous distribution across eukaryotes (**Fig. 1**). A C-terminal tyrosine is absent from β-tubulins in most eukaryotic groups examined, yet occurs sporadically in several distantly related lineages, with no obvious shared ecological or biological characteristic. Although limited by taxonomic representation in sequence databases, this distribution is compatible with either repeated losses of an ancestral β-tubulin tyrosine or independent gains in specific lineages. The presence of a terminal tyrosine also does not consistently correlate with that of known detyrosinases. For example, detyrosinated α-tubulin has been detected in *Drosophila* [49] and *Toxoplasma* [50] despite the absence of VASH or MATCAP/TMCP homologues (**Fig. S1**), indicating that additional tubulin carboxypeptidases remain unidentified.

Within Euglenozoa, the β-tubulin C-terminal tyrosine is absent from most bodonids but occurs in some prokinetoplastid, diplonemid and euglenoid lineages (**Fig. 1**). Its co-occurrence with VASH is also observed outside Euglenozoa in some heteroloboseans, including *Naegleria* (**Fig. S1**) where distinct tubulin isotypes form mitotic and flagellar MTs [51], and the C-terminal tyrosine is specifically associated with flagellar β-tubulin [52]. This association suggests that β-tubulin tyrosination may contribute to the specialization of distinct MT populations. This possibility is particularly relevant in trypanosomatids, where the conserved β-tubulin C-terminal tyrosine may help diversify MT populations despite a limited tubulin isotype repertoire. All trypanosomatids retain this terminal tyrosine together with a VASH homolog, coherent with a detyrosination machinery acting on both tubulin subunits [27,29].

The presence of both tyrosinated α- and β-tubulin raises the question of whether the two states are redundant or provide distinct information within the tubulin code. Our data show overlapping yet distinct distributions of tyrosinated α- and β-tubulin in *T. brucei* (**Fig. 2**). Tyrosinated β-tubulin persists across a broader range of MT structures than tyrosinated α-tubulin, consistent with less efficient VASH-mediated detyrosination of β-tubulin [29,30]. Within the axoneme, tyrosinated and detyrosinated β-tubulin preferentially localize to a single tubule of the outer doublets, revealing differential tyrosination within the A/B-tubule pair.

Thus, although α- and β-tubulin tyrosination largely coexist within the same MT populations, their distributions are not equivalent. Moreover, selectively shifting β-tubulin toward either the detyrosinated (**Fig. 3**) or tyrosinated (**Fig. 4**) state did not alter α-tubulin tyrosination, indicating that the tyrosination states of the two subunits can be independently regulated.

Despite independent regulation, the tyrosination states of both α- and β-tubulin converge on another major component of the tubulin code: glutamylation. Increased detyrosination of either β-tubulin in βΔ1 cells or α-tubulin in TTL KO cells increases glutamylation, whereas preventing β-tubulin detyrosination reduces it (**Fig. 5**). Glutamylation was lowest when both subunits remained tyrosinated in VASH KO cells, while restoring a mixed α/β tyrosination state in the βQ441D TTL KO double mutant returned glutamylation to near-parental levels. These genetic manipulations indicate that neither α-nor β-tubulin alone determines glutamylation. Instead, glutamylation reflects their combined tyrosination state. This extends the recently described crosstalk between detyrosination and glutamylation in trypanosomatids [28–30] to both subunits of the α/β heterodimer. Notably, constitutive β-tubulin detyrosination strongly increased GT335 signal but had a weaker effect on PolyE signal, suggesting a predominant effect on glutamylation acceptor sites rather than chain extension. Yet the β-monoE signal, which is expected to detect monoglutamylation of the β-tubulin E435 residue [44], remained unchanged under either constitutive detyrosination or tyrosination (**Fig. S5**). Recent mass-spectrometry analysis identified α-tubulin E445 and β-tubulin E438 as additional glutamylation sites, with longer glutamate chains occurring predominantly on α-tubulin [30]. Thus, the increased GT335 signal may reflect increased glutamylation at β-tubulin E438, α-tubulin E445, or both. Similar crosstalk between the two tubulin tails has been observed in *Tetrahymena*, where mutation of β-tubulin glycylation sites altered both glycylation and glutamylation of α-tubulin [53]. The presence of a regulatable C-terminal tyrosine on both subunits may therefore expand the control of glutamylation and its downstream MT functions.

Changes in β-tubulin tyrosination were accompanied by distinct morphological phenotypes. βΔ1 cells displayed a shorter free distal flagellum and a longer kinetoplast-to-nucleus distance, whereas βQ441D cells displayed the opposite (**Fig. 6**). In exponential phase, total flagellum length and new flagellum elongation were unaffected, arguing against impaired flagellum assembly and instead suggesting effects on MT organization or stability. Consistently, a similar shortening of the cell body has been reported following loss of tubulin detyrosination in stationary-phase *Leishmania* promastigotes, a phenotype rescued by paclitaxel treatment [29]. However, because tyrosination has little effect on purified tubulin dynamics *in vitro* [54,55], these morphological effects may involve MAPs and/or motors. Such a mechanism is consistent with the reported regulation of the flagellar kinesin-13.2 by tubulin detyrosination in *Leishmania* [29]. The opposing changes in free distal flagellum length in βΔ1 and βQ441D cells further suggest that the β-tubulin tyrosination state contributes to the spatial organization of the flagellum relative to the cell body.

This spatial organization may also extend to the axonemal MTs. In *T. brucei*, IFT trains become progressively restricted to specific axonemal doublets after entry through the transition zone, and tubulin PTMs such as glutamylation have been proposed to contribute to this track selection [56,57]. The spatial organization of β-tubulin tyrosination within the axoneme (**Fig. 2G-H**) suggests that this modification may also contribute to this specialization. In *Chlamydomonas*, tyrosinated and detyrosinated tubulin are enriched on the A- and B-tubules, respectively, which are preferentially used by anterograde and retrograde IFT trains [20]. Although we cannot assign these β-tubulin states to individual A- or B-tubules in *T. brucei*, normal new-flagellum elongation argues against a major IFT defect, while more subtle effects on IFT or axonemal motor activity remain possible.

The opposite motility phenotypes of the β-tubulin mutants further indicate that β-tubulin tyrosination controls flagellar function. Constitutive β-tubulin detyrosination reduced swimming speed and increased tumbling, whereas preventing detyrosination increased swimming speed and directional persistence while reducing tumbling (**Fig. 7**). These opposite effects parallel the opposite free flagellum lengths of the two mutants, although whether the two are causally related remains unknown. Since total flagellum length was unaffected, they could involve altered regulation of axonemal motors or waveform propagation. Notably, VASH KO cells showed no major motility defect, consistent with previous observations [27,28], indicating that simultaneous loss of α- and β-tubulin detyrosination does not reproduce the effects of selectively altering β-tubulin. This difference may reflect the distinct balance of α- and β-tubulin tyrosination and glutamylation in these backgrounds. At the collective level, the loss of organized SoMo in βΔ1 cells was particularly striking given their more moderate single-cell motility defect. Because SoMo integrates flagellar motility with environmental sensing and cAMP signaling [58], this phenotype suggests that the consequences of β-tubulin tyrosination may extend beyond motility to environmental responses relevant to parasite development within the tsetse fly.

More broadly, the functional consequences of β-tubulin tyrosination may depend on cellular and developmental context. Loss of tyrosinated β-tubulin in βΔ1 cells caused only a transient proliferation defect under standard culture conditions, suggesting that cells can adapt to a sustained change in the tubulin code (**Fig. S3G**). In contrast, a loss of tyrosinated α-tubulin in TTL KO cells did not cause any proliferation defect (**Fig. S4G**). However, perturbation of tubulin detyrosination has stronger consequences at specific developmental stages. In *Leishmania*, VASH deletion alters cell morphology and impairs amastigote proliferation, resulting in strongly reduced virulence in mice [29]. In *T. brucei*, VASH deletion also produces more severe morphological and proliferation effects in differentiation-competent pleomorphic cells [27]. The inability to recover our β-tubulin mutants in a pleomorphic background could similarly reflect a greater sensitivity of differentiation-competent parasites to perturbation of β-tubulin tyrosination. These findings raise the possibility that β-tubulin tyrosination becomes particularly important during parasite development, when changes in morphology, motility and environmental sensing are required for progression through different host environments. The motility defects observed in procyclic β-tubulin mutants could have even greater consequences in bloodstream forms, where motility is essential for survival [59].

Together, our findings establish β-tubulin tyrosination as an independently regulated component of the trypanosomatid tubulin code. Although rare across eukaryotes, the β-tubulin C-terminal tyrosine is conserved in trypanosomatids and occurs in several distantly related lineages. Its association with flagellar tubulin in *Naegleria* [51,52], together with its effects on flagellum positioning and motility in *T. brucei*, suggests a connection between β-tubulin tyrosination and flagellar MT function. In *T. brucei*, a C-terminal tyrosine on both α- and β-tubulin generates distinct but functionally connected states that converge on glutamylation. Thus, our study expands the canonical view of tubulin tyrosination and shows that the presence of a terminal tyrosine on α-tubulin, β-tubulin, or both can diversify MT regulation and function.

## Acknowledgements

We thank Diane McMahon-Pratt (Univ. of Yale) for the anti-PRF2 antibody, Mélanie Bonhivers (University of Bordeaux) for the mAb25 antibody and Jack Sunter (University of Edinburgh) for the pJ1339 plasmid. We acknowledge Elodie Jublanc from the imaging facility MRI, Montpellier and Chantal Cazevieille for her technical assistance in preparing electron microscopy samples and interpreting data concerning ultrastructural evaluation (University of Montpellier, INSERM U1298, Institute for Neurosciences of Montpellier, Montpellier). Special thanks to the EuPathDB team (https://eupathdb.org/eupathdb/). The work was funded by the following grants: FRM Equipe EQU202403018057 (K.R.), ANR-11-LABX-0024-01 (Y.S. and M.F.L.) and ANR-23-CE15-0022 (M.F.L.). Mass spectrometry experiments were carried out using the facilities of the Montpellier Proteomics Platform (PPM, BioCampus Montpellier), a member of the national Proteomics French Infrastructure (ProFI UAR 2048) supported by the French National Research Agency (ANR-24-INBS-0015, Investments for the future F2030). Work carried out in the Trypanosome Cell Biology Unit is funded by La Fondation pour la Recherche Médicale (EQU202203014654) and a French Government Investissement d’Avenir programme, Laboratoire d’Excellence “Integrative Biology of Emerging Infectious Diseases” (ANR-10-LABX-62-IBEID).

## Material and Methods

### Identification of orthologs

Orthologous groups were retrieved from two complementary databases: OrthoMCL v6 (https://legacy.orthomcl.org/orthomcl.l/app) and EggNOG 5.0 [35] (http://eggnog5.embl.de/#/app/home). OrthoMCL6 was preferred to OrthoMCL7 because of α- and β-tubulin cross-contamination within the ortholog groups, not present in OrthoMCL6. For each protein family of interest (α-tubulin; β-tubulin; VASH and MATCAP/TMCP), the corresponding ortholog group identifiers were extracted: OrthoMCL (OG6_100108; OG6_100137; OG6_105471; OG6_107867) and EggNOG (KOG1376; KOG1375; 2QPPX; 2QQGI).

For some genes absent from both ortholog databases, additional homologs were identified using BLASTp searches against NCBI nr and published literature, including the *Chlamydomonas reinhardtii* VASH2-SVBP sequence [20] and additional Euglenozoan proteome and genome [60–62]. Protein FASTA entries of tubulins were screened to identify sequences that end with a tyrosine residue (Y) using a custom Python script with Biopython (SeqIO). Full length matching sequences were retained and pooled into a single curated dataset including supergroup, subclade, genus and binary presence/absence values for each ortholog. Heatmap displaying the presence/absence patterns of orthologous proteins were generated using a custom Python script with pandas and matplotlib and exported as SVG files.

### Cell culture

Procyclic 427 *T. brucei* cells were cultivated in SDM-79 (HyClone Laboratories) supplemented with 10% heat-inactivated fetal bovine serum (Gibco) and 7 µg/L hemin at 27°C.

### Generation of *T. brucei* transgenic cell lines

To delete the VASH (Tb927.11.1060) and TTL (Tb927.2.5250) genes using a PCR-based CRISPR-Cas9 strategy [63], procyclic *T. brucei* 427 cells were transfected with the pJ1339 plasmid, derived from pJ1173 [64], which carries a puromycin resistance marker, the Tet repressor, T7 RNA polymerase, and Cas9. Deletion of the tyrosine codon of β-tubulin was performed using the marker free CRISPR-Cas9 as previously described [42]. Briefly, crRNA guides were ordered on Integrative DNA Technologies (IDT) and duplexed with tracrRNA. The duplex was incubated with the spCas9 (IDT, SpCas9 nuclease V3) 20 minutes before transfection. Transfection was performed by washing 3.10^6^ cells in transfection buffer (187,5 µM CaCL2; 83 mM Na2HPO4; 29 mM NaH2PO4; 6,22 mM KCl; 62,2 mM HEPES pH 7,3) and resuspended in transfection buffer supplemented with the spCas9 mixture and a PCR-based donor DNA. Nucleofection was performed on the Amaxa Nucleofector 2b (Lonza) using program X-001. Cells were then cloned by serial dilutions in a 96 well plate to isolate edited cells.

### Ultrastructure expansion microscopy

*T. brucei* cells from parental, βΔ1 and VASH KO were loaded on 12-mm coverslips in a 24-well plate and cells left to adhere 10 minutes. Cells were then covered with 1 mL of fresh fixation solution (4% formaldehyde; 4% Acrylamide in PBS) and incubated for 3 hours at 37°C. For the gelation step, coverslips were deposited in a gelation chamber as described in Louvel et al., 2023 [65]. 50µL of activated Monomer solution was added (19% Sodium acrylate (AK Scientific); 10% Acrylamide (Euromedex); 0.1% DHEBA (N,N’-(1,2-Dihydroxyethylene)bisacrylamide; Sigma-Aldrich), 0.5% TEMED (Euromedex), 0.5% Ammonium persulfate (Euromedex) in PBS) for 15 min on ice then transferred at 37°C and incubated for 1 hour in a moist chamber. The coverslips were then transferred in 6-well plate in 1 ml of denaturation solution (200 mM Sodium Dodecyl Sulfate; 200 mM NaCl; 50 mM Tris pH 6,8) with agitation at RT to detach the gel from the coverslip, then moved into a 1.5 ml Eppendorf centrifuge tube filled with 1mL of denaturation solution and incubated at 95°C for 90 min. Gels were expanded in 50 ml of deionized water three times then shrinked in 50 ml of PBS for 15 minutes three times. Next, gel portions were incubated with primary antibodies diluted in blocking solution (PBS 2% BSA) overnight at 37°C with agitation (rabbit anti-αΔ1-tubulin, 1:1000 dilution; rabbit anti-βΔ1-tubulin, 1:1000 dilution; rabbit anti-β-tyr-tubulin, 1:1000 dilution; rat YL1/2, 1:750 dilution; mouse GT335, 1:1000 dilution; rabbit PolyE, 1:1000 dilution; mouse anti-β-tubulin, 1:200 dilution). After three washes in blocking solution, gels were incubated with secondary antibodies conjugated to Alexa Fluor 488 (1:400 dilution) or Alexa Fluor 594 (1:400) (Thermo Scientific) diluted in blocking solution with 57 µM Hoechst 33342 (Thermo Scientific) for 3 hours in the dark at 37°C with agitation. After three PBS washes, gels were expanded with three washes of deionized water. Acquisition was performed on a confocal microscope (Zeiss LSM880) equipped with an Airyscan detector with a 63X oil objective (NA = 1.4) using Zen Black software (Zeiss, Intelligent Imaging Innovations). Z-stacks were processed to obtain maximum intensity projections.

### Iterative ultrastructure expansion microscopy

Iterative expansion was performed as described by Louvel et al (Nat. Comm 2023) [65]. Briefly, *T. brucei* 427 cells were washed twice in 1x PBS and 10^7^ cells in 50µl PBS were adhered for 10 minutes on 12 mm coverslips coated with poly-L-Lysine. Coverslips were transferred in 1 ml post fixation solution (PFS) containing 1.4% paraformaldehyde and 2% acrylamide in 1× PBS for 3 h at 37°C. For the gelation, gelation chambers were made as described (Louvel et al, 2023) [65] and placed on ice. Sample coverslips were transferred to the chambers and covered with monomer solution (10% acrylamide, 19% sodium acrylate, 0.1% DHEBA, 0.25% tetramethyl ethylenediamine (TEMED) and 0.25% ammonium persulfate). Samples were incubated for 15 minutes on ice, then polymerization was completed at 37°C for 45 minutes in a humidified chamber. The coverslips were then removed from the gelation chamber, transferred in Eppendorf tubes containing 1mL denaturation buffer (200 mM SDS, 200 mM NaCl, 50 mM Tris base, pH 6.8) and incubated for 1hr at 85°C. Following denaturation, gels were washed 3 times 30-min in deionized water to initiate expansion. Expanded gels were subsequently shrinked in PBS then co-immunostained overnight at 37°C using mouse IgG2b 6.11.B1 anti-acetylated antibody (Piperno et al. J Cell Biol 1987) [66] and rabbit anti-βΔ1 or anti-βtyr . All antibodies were diluted 1/500 in 2% BSA in PBS. Following primary antibody incubation, gels were washed three times with PBS-Tween 20 for 15 minutes, then incubated for 3 hours in the dark with secondary antibodies Abberior STAR RED, goat anti-rabbit IgG and Abberior STAR ORANGE, goat anti-mouse IgG diluted 1/200 in 1x PBS containing 2% BSA. After secondary antibody incubation, gels were washed two times with PBS-Tween 20 before staining with DAPI (2µg/mL in PBS) and three washes in deionized water for 30 minutes each. 1.5×1.5 cm gel sample were re-embedded in a second gel for further expansion. For re-embedding, gels were placed in 6-well plates containing 1.5 mL of 2 neutral monomer mix (10% acrylamide, 0.05% DHEBA, 0.05% APS/TEMED) and incubated on ice with gentle agitation 3 times for 10mn each, lid open. Gels were subsequently mounted on microscope slides, excess monomer was blotted away using Kimwipes, and a 22 × 22 mm coverslip was applied. Polymerization was carried out in a humidified chamber at 37°C for 1 h. Following embedding, the gels were treated with an PFS at 37°C for 3 h under shaking, washed twice with PBS, and then three times 10 minutes in a final expansion monomer solution (10% acrylamide, 19% sodium acrylate, 0.1% N,N’-methylene-bis-acrylamide, 0.05% APS/TEMED) on ice with agitation. After removing excess solution, gels were covered with coverslips, and polymerization was completed at 37°C for 1 h in a humidified environment. Dissolution of neutral and first gel was performed following gels treatment with 200 mM NaOH for 1 h at room temperature, followed by multiple washes in PBS (10min each) until neutral pH was achieved. Full expansion was accomplished through successive incubations in deionized water.

Confocal and STED images were acquired on a STED expert line microscope (Abberior Instruments) using a 100×/1.4 NA oil objective (Olympus) and Avalanche Photo-diodes (single-point photon counting modules) detector (Excelitas). The two-color STED images were obtained in line-interleaved mode, switching on one excitation laser line at a time, 561 nm for Abberior StarOrange and 640 nm for Abberior StarRed. The laser line for STED was 775 nm, pulsed. Pinhole size was set to 0.8 Airy unit. In confocal mode, pixel size was 50nm and elementary pixel dwell-time was 3 μs with 40 line averaging. Step size between images for 3D acquisitions was set to 1 µm. In STED mode, pixel size was 20nm and elementary pixel dwell-time was 1.5µs with 160 line averaging for STAR Orange and 80 for STAR Red.

### Quantitative PCR

Genomic DNAs were extracted using the NucleoSpin® Blood QuickPure kit (Macherey-Nagel) according to the manufacturer’s instructions. Quantitative PCR (qPCR) assays were performed on a LightCycler® 480 Real-Time PCR System (Roche) using the LightCycler® 480 SYBR Green I Master mix (Roche). The amplification program consisted of an initial denaturation step at 95°C for 5 min, followed by 44 cycles of denaturation at 95°C for 10 s, annealing at 58°C for 10 s, and extension at 72°C for 10 s. The relative abundance of each tubulin gene was normalized to the housekeeping gene GPI8, used as an internal reference. For each primer pair, three consecutive 10-fold dilutions were analyzed: 1, 0.1 and 0.01 ng/µL. Each dilution was analyzed in four technical replicates, and the experiment was performed in three biological replicates.

### Quantification and statistical analysis

Quantitative analyses were performed using non-parametric statistical testing. For all comparisons involving more than two groups, a Kruskal–Wallis test was applied followed by Dunn’s post-hoc test for multiple comparisons. At least three independent biological replicates were included in each experiment. Morphometric measurements were performed blindly by an independent operator to avoid selection or measurement bias. For morphometric analyses, statistics were performed on the mean of each independent experiment, and mutants were compared with the parental cell line within each growth phase using a repeated-measures one-way ANOVA followed by Dunnett’s multiple comparisons test. Distance measurements were carried out using Zen Blue software (Zeiss), and all statistical analyses and graphical representations were generated in GraphPad Prism 8 and 11.

### Western blot

For western blotting analysis, *T. brucei* whole-cell or cytoskeleton extracts were separated by SDS-PAGE and transferred onto PVDF membranes (Hybond-P, Amersham). Membranes were incubated overnight at 4 °C with primary antibodies diluted in PBS 1X 0,1% tween 2% BSA. After washing, the membranes were incubated with HRP-conjugated secondary antibodies for 1 hour at room temperature and revealed with Clarity ECL substrate (Biorad). Membrane imaging is performed using Chemidoc^TM^ MP Imaging System (BioRad).

### Immunofluorescence assay

Procyclic cells were harvested by centrifugation, washed twice in PBS and settled on microscopic slides for 10 minutes and fixed with paraformaldehyde (PFA) for 4 minutes. To extract cytoskeleton, settled cells were treated with 0,25% Nonidet P-40 in a PIPES buffer (0,1M PIPES pH 6,9; 1mM MgCl2) for 1 minute and washed with PIPES buffer before PFA fixation. After three PBS washes, cells were blocked in 2% BSA for 1 hour. Incubation with primary antibodies was performed overnight at 4°C or 3 hours at room temperature in blocking solution (rabbit anti-βΔ1-tubulin, 1:3000 dilution; rabbit anti-β-tyr-tubulin, 1:3000 dilution; rat YL1/2, 1:750 dilution; mouse mAb25, 1:5 dilution; rabbit anti-FLAM8, 1:500 dilution; mouse anti-PFR2 L8C4, 1:500; rabbit anti-β-tubulin AA344, 1:1000). Slides were washed three times with blocking solution before incubation with secondary antibodies for 1 hour (Thermo Fisher anti-IgG specific secondary antibodies conjugated to Alexa Fluor 488 or 594 at a dilution of 1:2000). Slides were then washed twice in PBS and DNA was stained using Hoechst 33342 and mounted in Slowfade gold medium (Invitrogen). Imaging was performed using Zen Blue 3.6 (Zeiss) software, on a Zeiss imager Z1 microscope using a Hamamatsu digital camera C11440 with objective 100X NA 1.4.

### Motility assay

Cells were taken during early exponential growth at a density of 1-2×10^6^ cells/mL. 8µL of cell culture were deposited in a FastRead 102® (BVS100, Biosigma) cell counting slide. Cells were imaged in brightfield for 60 seconds at a frequency of five images per second using a 20X/0,5 Ph2 Neofluar objective. Motility parameters were measured using the Fiji plugin “Combined Particle Tracking” with Gaussian blur radius set to 15px and Noise tolerance to 150.

### Social motility assay

The protocol was adapted from [67]. Briefly, 45 mL of SDM79 supplemented with 10% iFBS and 20mM glycerol were pre-warmed to 42°C. 5 mL of low-melting point agarose (Gibco BRL, 4% w/v in water) was added to the pre-warmed medium. 12 mL of the resulting 0,4% agarose media were immediately poured into 90 mm petri dishes. Plates were air-dried for 1 hour in a laminar flow cabinet. *T. brucei* procyclic cells in early exponential phase (1-2×10^6^ cells/mL) were centrifuged and resuspended in 100 µL of residual medium at a density of 1,2 ×10^5^ cells/µL. Five µL were spotted on the surface of the plates and allowed to absorb for 10 minutes. Plates were sealed with parafilm and incubated at 27°C. SoMo communities were imaged in a Chemidoc^TM^ MP Imaging System (BioRad) in colorimetric mode with plates opened.

### Nanopore sequencing

High molecular weight DNA was extracted from 1.10^8^ cells using the kit Monarch® HMW DNA extraction kit for cells & blood (T3050L, NEB). DNA quality and concentration was analysed by Nanodrop (Thermo Scientific) and DeNovix DS-11 FX+ (DeNovix), respectively. Libraries were prepared following the manufacturer’s protocol using the Rapid Sequencing Kit V14 -gDNA (SQK-RAD114). Libraries were then loaded on a MinION device and reads were acquired on MinKNOW software. Reads were then aligned against *T. brucei* genome (*Trypanosoma brucei* Lister strain 427 2018) and viewed on IGV (Integrative Genomics Viewer).

### Flagellar extraction

Cells were washed with PBS and subjected to cytoskeletal extraction in 0.1 M PIPES buffer containing 1 mM MgCl₂ and 1% NP-40. Flagella were then extracted by incubating the cytoskeletal fraction for 30 min at room temperature in the same buffer supplemented with 1 M KCl. Extraction efficiency was assessed by examining an aliquot by light microscopy. The recovered flagella were washed with 0.1 M PIPES containing 1 mM MgCl₂, resuspended in Laemmli sample buffer, and heat-denatured at 95°C for 10 min.

### Mass spectrometry

Samples were prepared and analyzed as previously described [8]. Briefly, after separation by SDS–polyacrylamide gel electrophoresis, tubulin regions were isolated and proteins were in-gel digested using trypsin (1 μg per band; Gold; Promega). Peptides were then analyzed online by nanoflow HPLC–nanoelectrospray ionization using a Q Exactive HFX mass spectrometer (Thermo Fisher Scientific) coupled to a nano-LC system (U3000-RSLC, Thermo Fisher Scientific). A gradient consisting of 0 to 40% B in A for 60 min (A: 0.1% formic acid, 2% acetonitrile in water and B: 0.1% formic acid in 80% acetonitrile) at 300 nl/min was used to elute peptides from the capillary reverse-phase column (0.075 mm by 250 mm, Pepmap, Dionex). Data were acquired using the Xcalibur software (version 4.7.69.37). A cycle of one full-scan mass spectrum (350 to 1500 m/z) at a resolution of 120,000 (at 200 m/z), followed by 20 data-dependent MS/MS spectra (at a resolution of 30,000; isolation window, 1.6 m/z) was repeated continuously throughout the nano-LC separation. Raw data analysis was performed using the MaxQuant software (version 2.0.3.0) with standard settings. Used database consist of Trypanosoma brucei Reference proteome entries from Uniprot (www.uniprot.org, V2026_02, UP000008524), and tubulin sequences without Cterminal trysosine and 250 classical contaminants (MaxQuant contaminant database). Relative abundance of peptide was estimated using Skyline 21.2.

### Scanning electron microscopy

Stationary phase *T. brucei* cells washed in PBS were fixed with 2.5% glutaraldehyde in PHEM buffer, pH 7.2 for an hour at room temperature, followed by washing in PHEM buffer. Fixed samples were dehydrated using a graded ethanol series (30-100%), followed by 10 minutes in graded Ethanol - Hexamethyldisilazane. And then Hexamethyldisilazane alone. Subsequently, the samples were sputter coated with an approximative 10nm thick gold film and then examined under a scanning electron microscope (Hitachi S4000, at CoMET, Intitut des Neurosciences Montpellier, INSERM, Université de Montpellier, Montpellier France,) using a lens detector with an acceleration voltage of 10KV at calibrated magnifications.

---

**Figure S1. (related to Figure 1).**
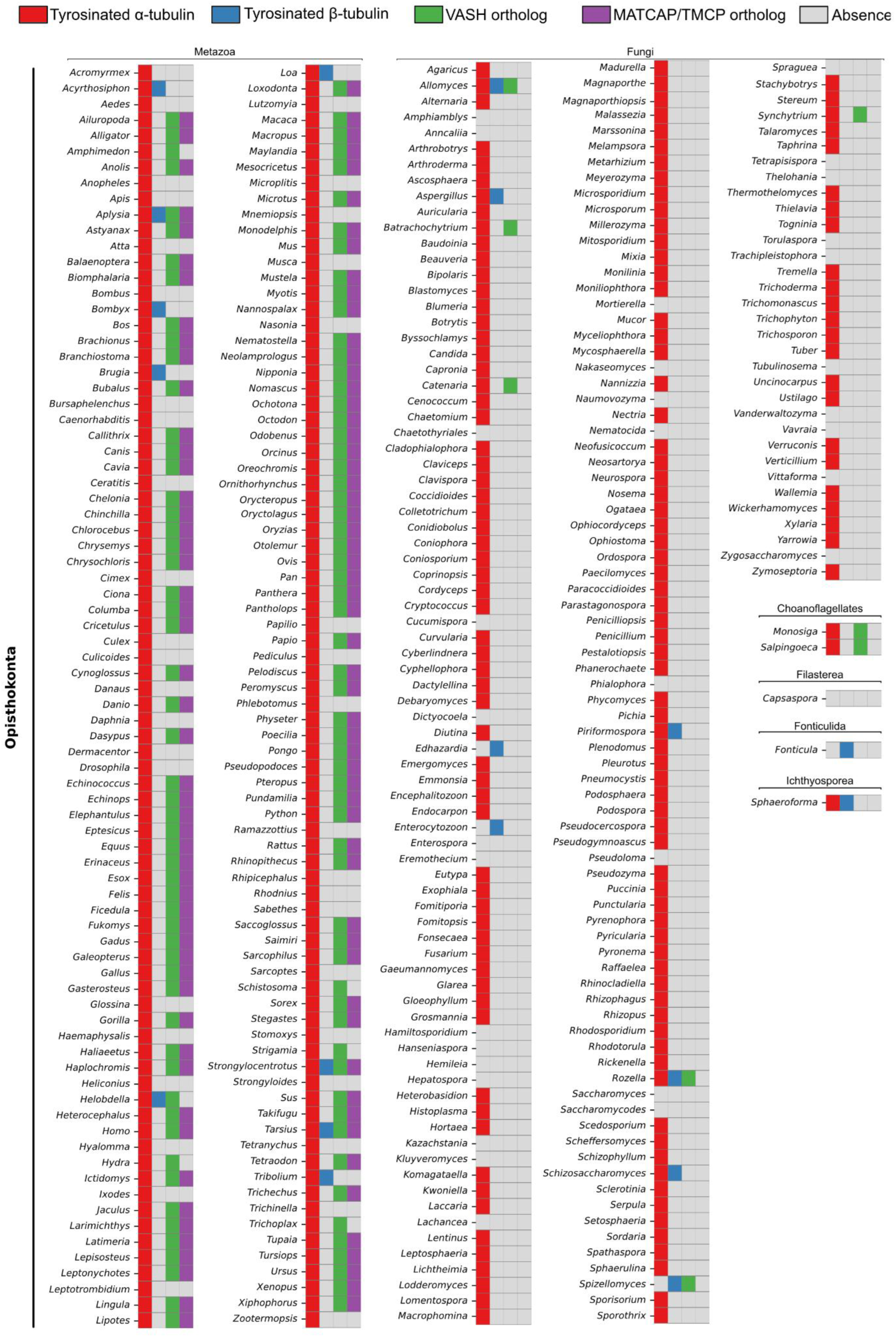
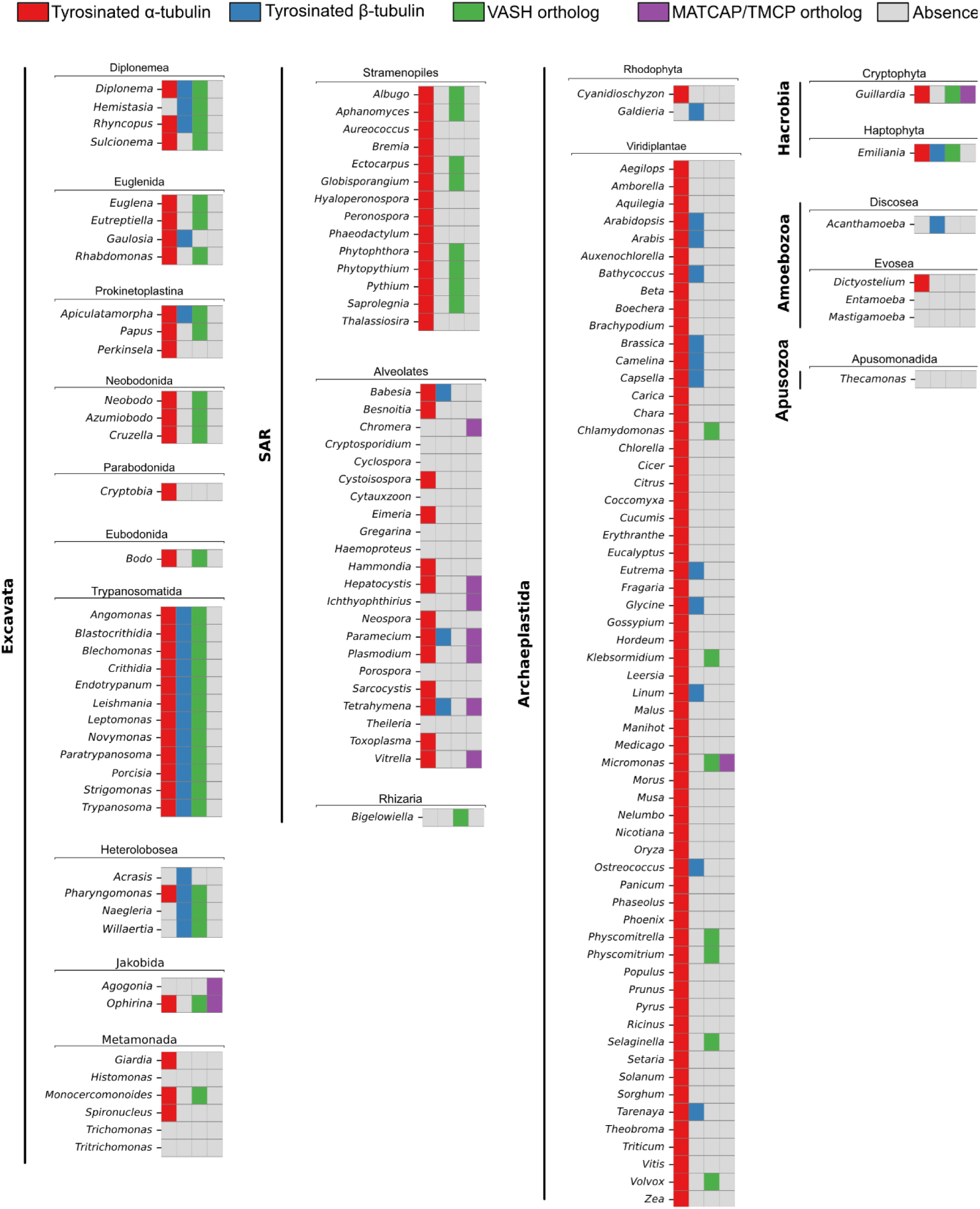
β-tubulin tyrosination is an evolutionarily conserved feature of trypanosomatids. Heatmap compiling the presence or absence of C-terminus tyrosine motifs and detyrosinase across eucaryotic genus. The presence or absence of protein motif and enzyme was determined from each OrthoMCL6 and EggNOG groups corresponding to α-tubulin (OG6_100108 and KOG1376), β-tubulin (OG6_100137 and KOG1375), VASH (OG6_105471 and 2QPPX) and MATCAP/TMCP (OG6_107867 and 2QQGI) proteins. From left to right, colored rectangles correspond to the presence of tyrosinated α-tubulin (red), tyrosinated β-tubulin (blue), VASH ortholog (green), and MATCAP ortholog (purple). Grey indicates absence of tyrosine motifs or enzyme.

**Figure S2. (related to Figure 2).**
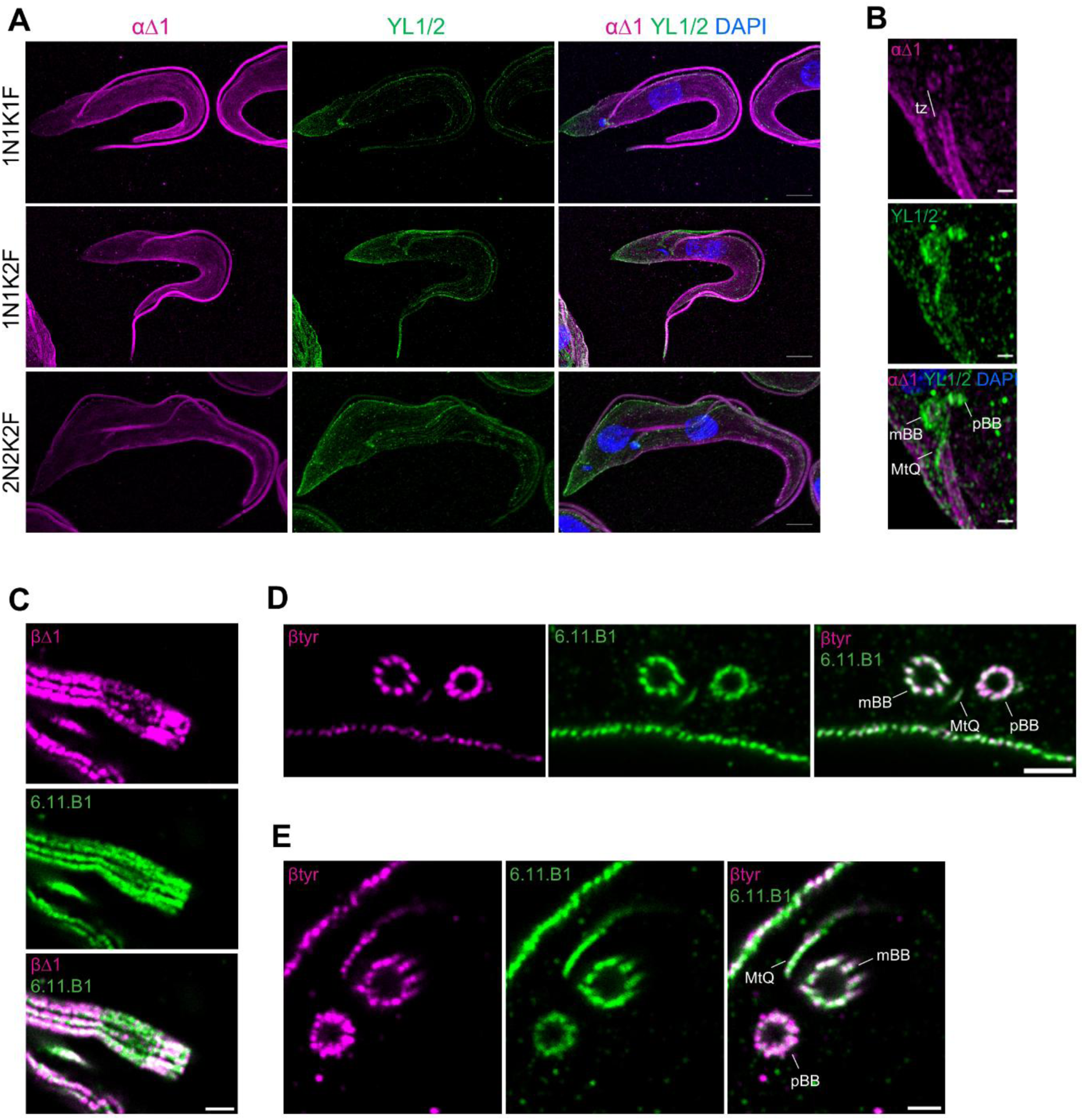
Distinct subcellular distribution of α- and β-tubulin tyrosination in *T. brucei*. (**A-B**) UexM analysis of tyrosinated and detyrosinated α-tubulin in *T. brucei* parental procyclics. Maximum intensity projections of z-stack confocal images are presented. The αΔ1 antibody recognizes detyrosinated α-tubulin. Scale bar: 10µm (A) or 1µm (B). (**C**) iUexM analysis of detyrosinated β-tubulin in *T. brucei* parental procyclics. Maximum intensity projections of z-stack confocal images are presented. The 6.11.B1 antibody recognizes acetylated α-tubulin. Scale bar: 2µm. (**D-E**) iUexM analysis of tyrosinated β-tubulin in *T. brucei* parental procyclics in axonemal cross sections. Single Z-stack are presented. Scale bar: 2µm.

**Figure S3. (related to Figure 3).**
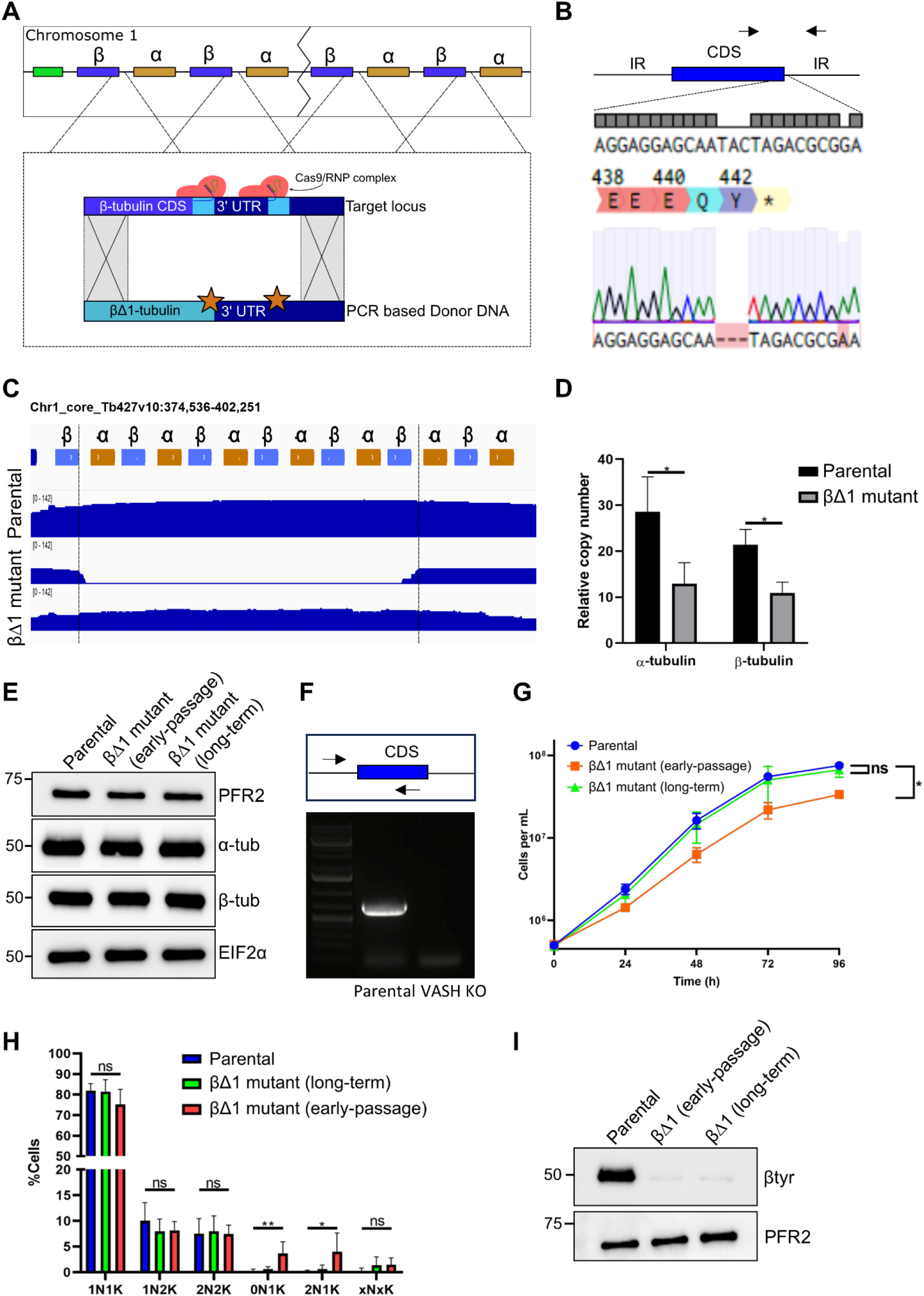
Engineering a constitutively detyrosinated β-tubulin state. (**A**) Schematic representation of the CRISPR/Cas9 strategy to generate the βΔ1 mutant. (**B**) Sanger sequencing of the β-tubulin locus from βΔ1 mutant genomic DNA aligned to the reference genome, confirming the absence of the tyrosine codon. (**C**) Nanopore sequencing analysis of the α- and β-tubulin tandem array located on chromosome 1 performed on genomic DNA from parental and long-term cultured βΔ1 mutant cell lines. (**D**) qPCR analysis of α- and β-tubulin copy number performed on genomic DNA from parental and long-term cultured βΔ1 mutant cell lines. GPI8 control gene was used for normalization (n= 3 from three independent DNA extractions). (**E**) Immunoblot analysis of α- and β-tubulin abundance in whole cell extracts from parental, long-term and early-cultured βΔ1 mutant. PFR2 and EIF2α are used as loading controls. (**F**) Diagnostic PCR of the VASH KO cell line. Schematic representation of TbVASH gene locus (blue) and primers (black arrows) used to confirm the loss of wild-type allele. Parental cell line is used as a positive control. (**G**) Growth curve (log scale) of *T. brucei* procyclic cells from parental, long-term and early-cultured βΔ1 mutant cell lines cultured over 4 days. (**H**) Quantification of Nuclei (N) and kinetoplasts (K) in *T. brucei* parental, long-term and early-cultured βΔ1 mutant cell lines during exponential growth. (n = 4; error bars represent SD). (**I**) Immunoblot analysis of β-tubulin tyrosination status in whole cell extracts from parental, long-term and early-cultured βΔ1 mutant cell lines. PFR2 is used as loading control.

**Figure S4. (related to Figure 4).**
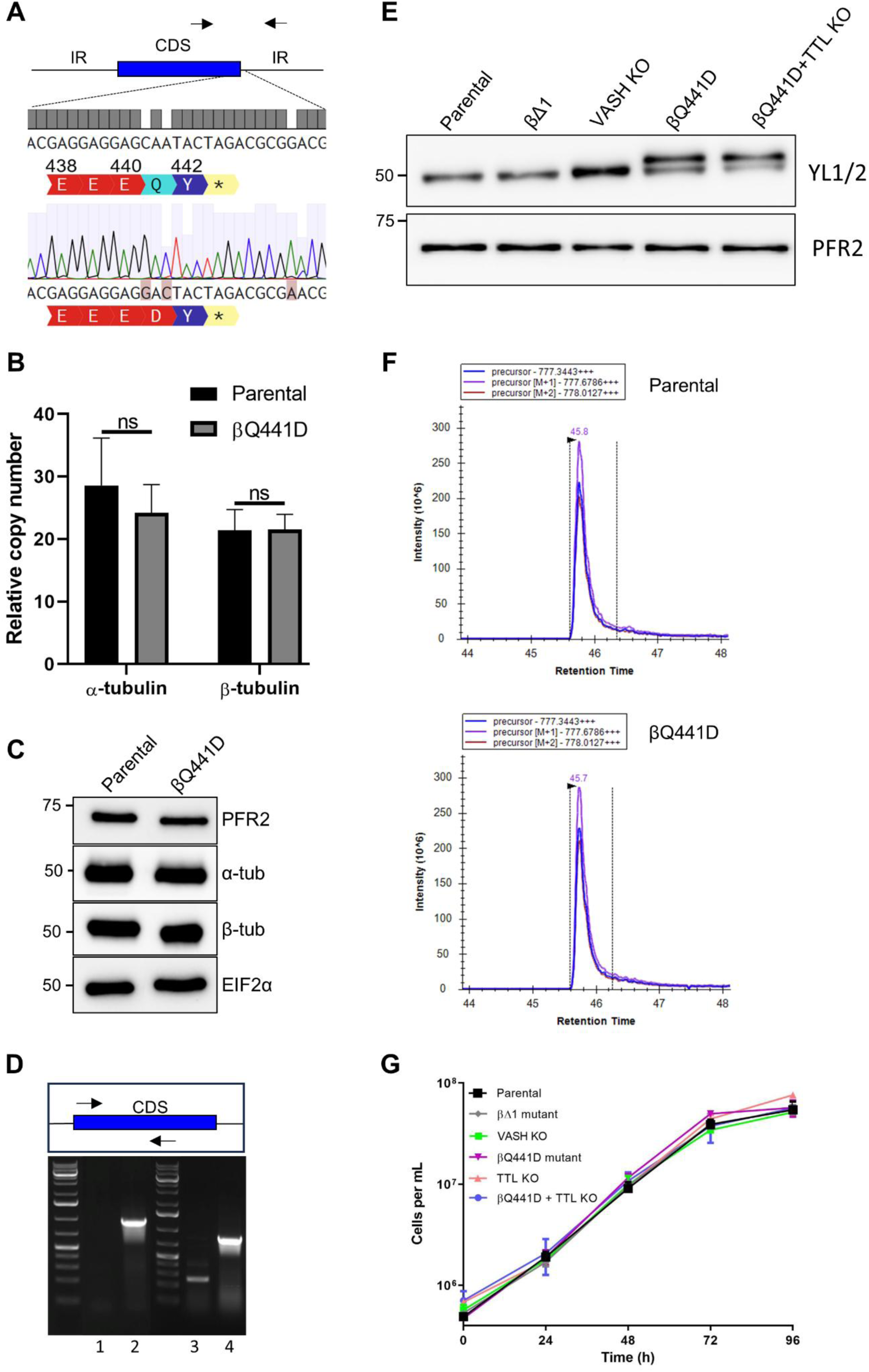
Engineering selective inhibition of β-tubulin detyrosination. (**A**) Sanger sequencing results of the βQ441D mutant genomic DNA mapped against the reference genome validating the complete edition of the glutamine codon to an aspartate. (**B**) qPCR analysis of α- and β-tubulin copy number performed on genomic DNA from parental and βQ441D mutant cell lines. GPI8 control gene was used for normalization (n= 3 from three independent DNA extractions). (**C**) Immunoblot analysis of α- and β-tubulin abundance in whole cell extracts from parental and βQ441D mutant cell lines. PFR2 and EIF2α are used as loading controls. (**D**) Diagnostic PCR of the TTL KO in parental and βQ441D backgrounds. Schematic representation of TbTTL gene locus (blue) and primers (black arrows) used to confirm the loss of wild-type allele. Parental cell line is used as a positive control. 1 : TTL KO in parental background; 2: Parental control cell line; 3 : TTL KO in βQ441D background; 4 : Parental control cell line. (**E**) Immunoblot analysis of YL1/2 reactivity in whole cell extracts from parental, βΔ1 mutant, VASH KO, βQ441D mutant and βQ441D TTL KO. PFR2 is used as a loading control. (**F**) Mass spectrometry analysis of internal α-tubulin peptides from parental and βQ441D mutant cell lines. Representative extracted ion chromatograms corresponding to α-tubulin peptides located upstream of the C-terminal tail are shown, highlighting the equivalent quantity between the two cell lines. (**G**) Growth curve (log scale) of *T. brucei* procyclic cells from parental, βΔ1 mutant, VASH KO, βQ441D mutant, TTL KO early and late cultures, and βQ441D TTL KO cell lines cultured over 4 days.

**Figure S5. (related to Figure 5).**
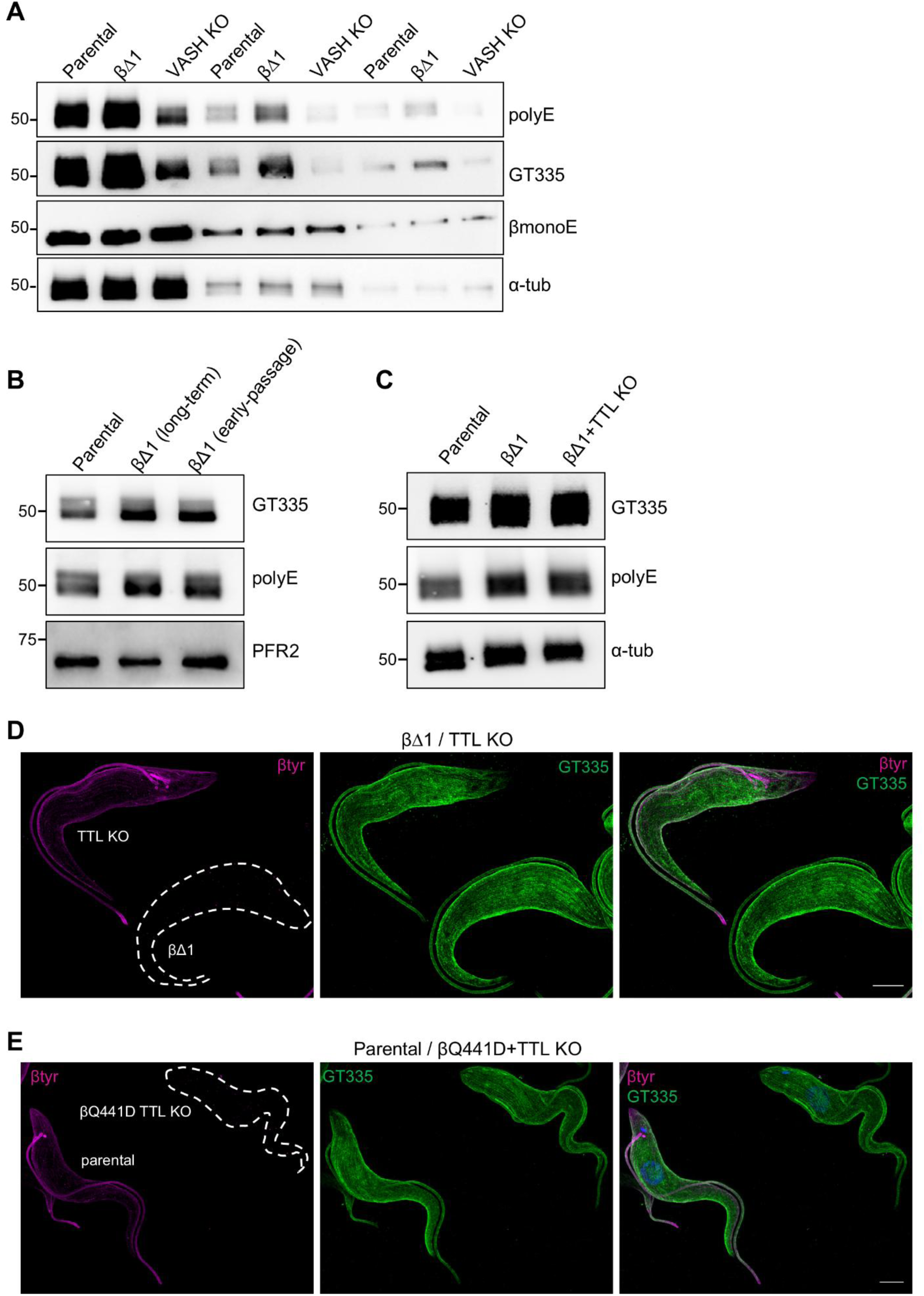
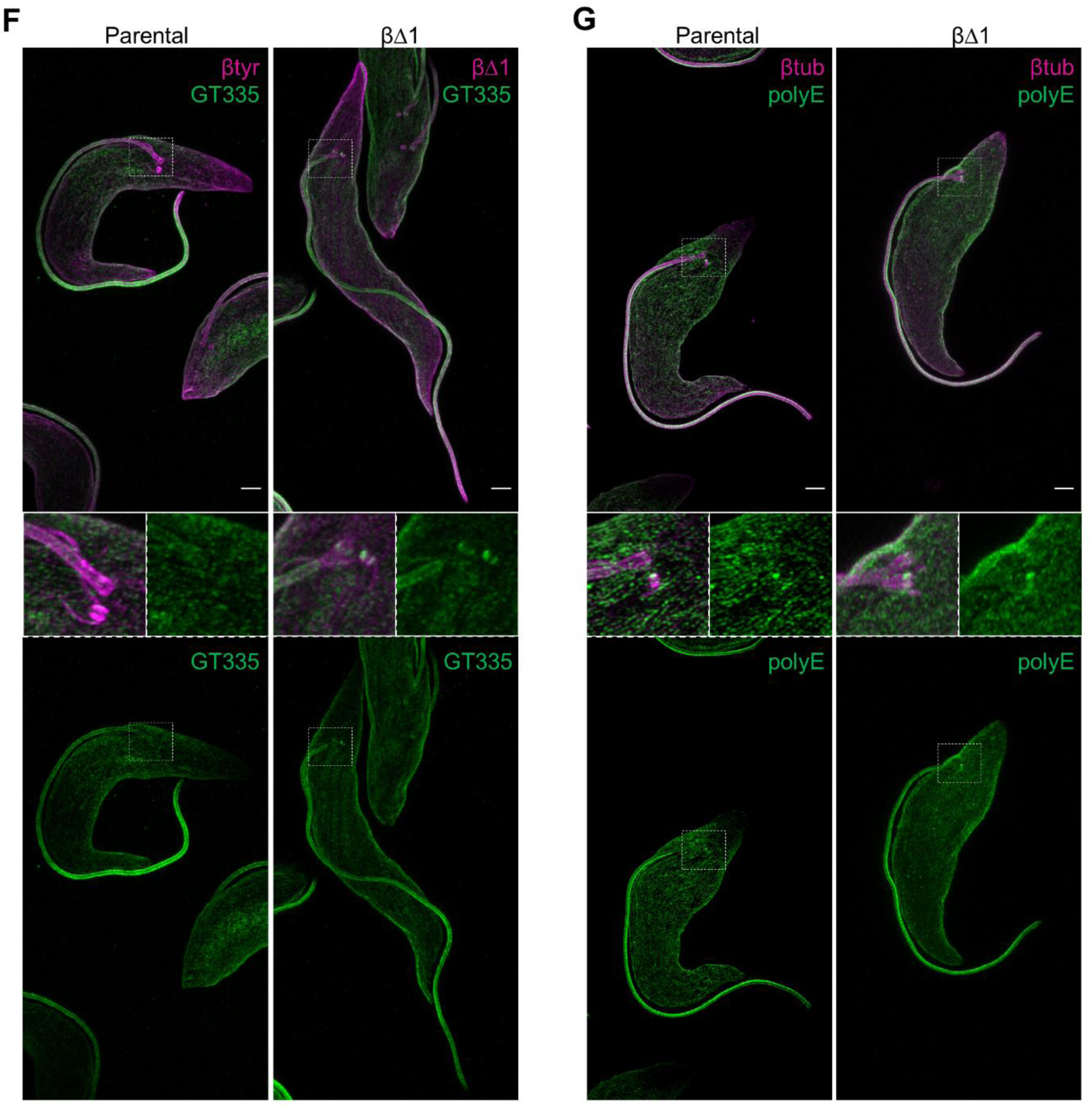
β-tubulin tyrosination independently regulates microtubule glutamylation. (**A**) Immunoblot analysis of glutamylation levels in threefold serial dilutions of whole cell extracts from parental, βΔ1 mutant and VASH KO cell lines. The βmonoE antibody recognizes the branching point glutamate of β-tubulin. α-tubulin is used as a loading control. (**B**) Immunoblot analysis of glutamylation levels in long-term and early-cultured βΔ1 mutant populations. PFR2 is used as a loading control. (**C**) Immunoblot analysis of glutamylation levels in whole cell extracts from parental, βΔ1 and βΔ1 TTL KO mutant cell lines. α-tubulin is used as a loading control. (**D-E**) UexM analysis of microtubule glutamylation of βΔ1 mutant and TTL KO cell lines (D) or parental and βQ441D TTL KO mutant cell lines (E) embedded in the same gel. Cells are labelled with GT335 and β-tyr antibodies. Maximum intensity projections of z-stack confocal images are presented. Scale bar : 10µm. (**F**) U-exM analysis of parental and βΔ1 mutant stained with GT335 and co-labeled with either β-tyr or βΔ1 antibodies as basal body markers. Parental and βΔ1 mutant cell lines were embedded and processed in separate gels. Maximum intensity projections of z-stack images are presented. Scale bar: 5µm. (**G**) U-exM analysis of parental and βΔ1 mutant stained with PolyE and co-labeled with β-tubulin. Parental and βΔ1 mutant cell lines were embedded and processed in separate gels. Maximum intensity projections of z-stack images are presented. Scale bar: 5µm.

**Figure S6. (related to Figure 6).**
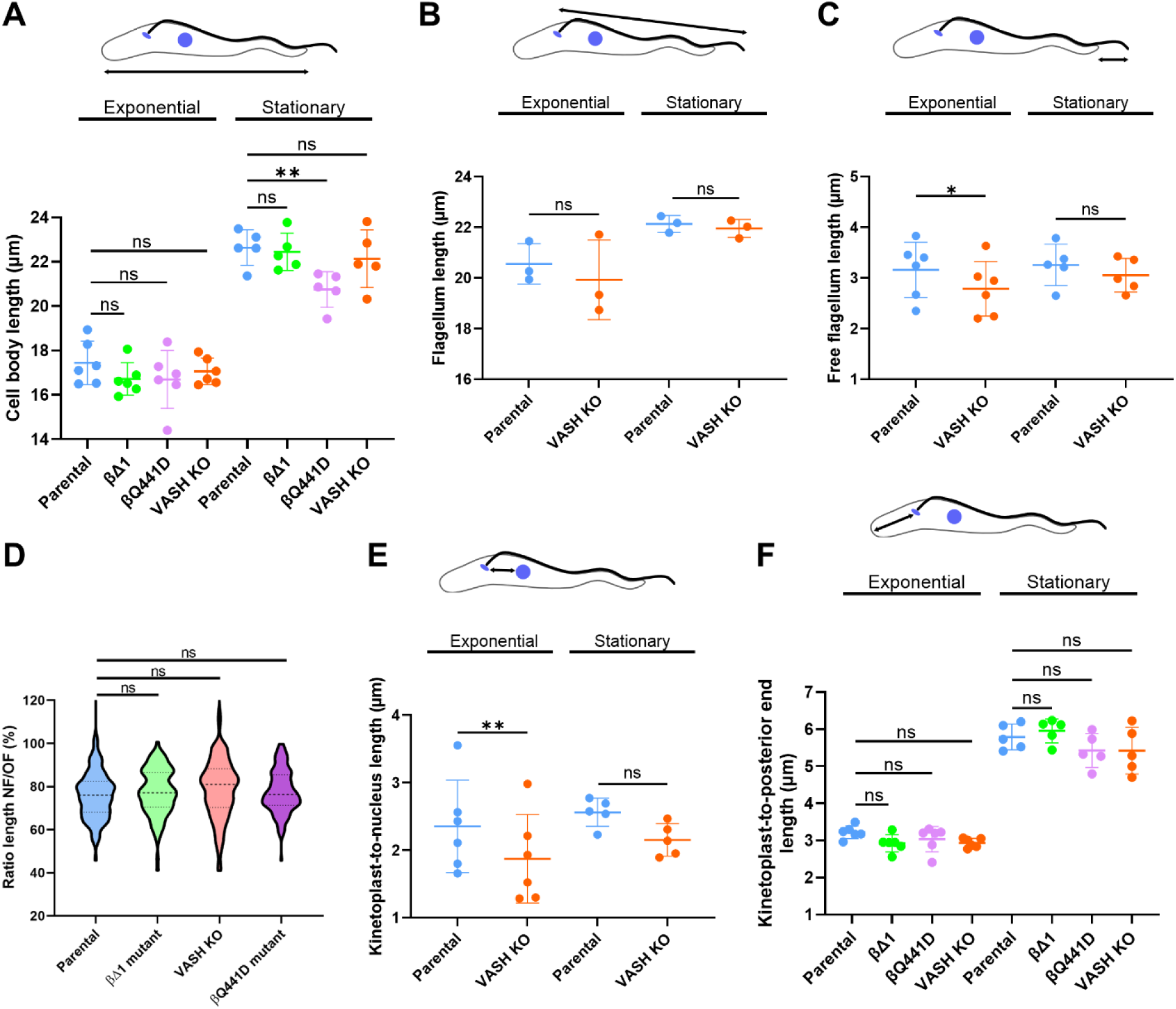
β-tubulin tyrosination impacts parasite morphology. (**A**) Measurements of cell body length from parental, βΔ1 mutant, βQ441D mutant and VASH KO cell lines cultured in exponential and stationary phase (six and five independent experiments, respectively). (**B-C**) Measurements of total flagellum length (B), three independent experiments per growth phase) and free flagellum length (C), six and five independent experiments in exponential and stationary phase, respectively) from parental and VASH KO cell lines. (**D**) Ratio between the length of the new (NF) and old flagella (OF) in 2K2N cells displaying FLAM8 at the tip of both flagella, from parental, βΔ1 mutant, βQ441D mutant and VASH KO cell lines cultured in exponential phase. (**E**) Measurements of kinetoplast-to-nucleus distance from parental and VASH KO cell lines cultured in exponential and stationary phase (six and five independent experiments, respectively). (**F**) Measurements of kinetoplast-to-posterior end distance from parental, βΔ1 mutant, βQ441D mutant and VASH KO cell lines cultured in exponential and stationary phase (six and five independent experiments, respectively). (A-C, E-F) Each point represents the mean of one independent experiment (n=50 cells per experiment) and bars show the mean ± SD. Flagellum length was measured on mAb25-labelled cells; other parameters were measured on mAb25- or PFR2-labelled cells. Cells were labelled with β-tubulin for the cell body and Hoechst 33342. The schematic above each graph indicates the parameter measured (highlighted with the arrow). Each cell line was compared with the parental cell line within the same growth phase using a repeated-measures one-way ANOVA including all four cell lines, followed by Dunnett’s multiple comparisons test (ns, not significant; *P<0.05; **P<0.01).

**Table S1.**
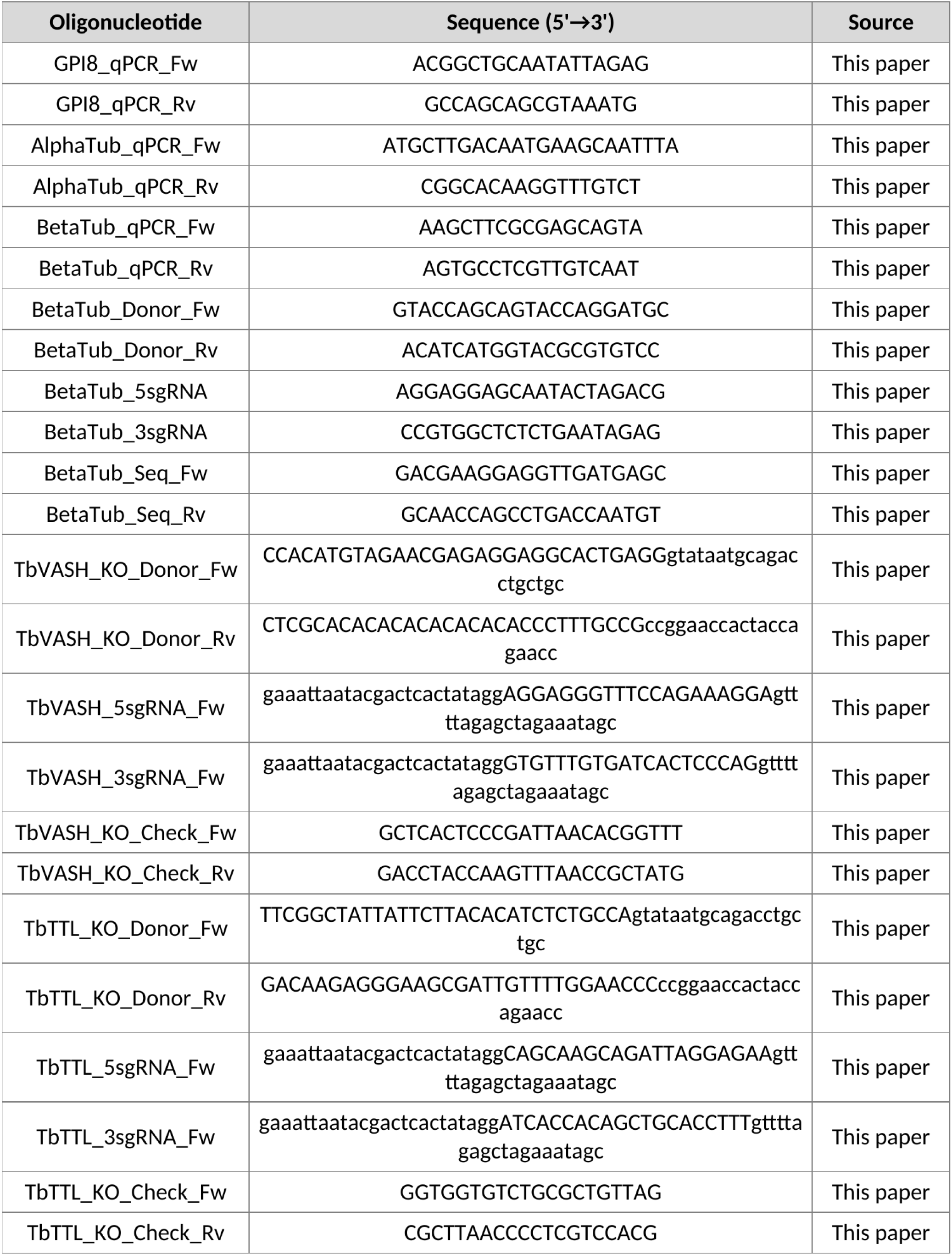
Oligonucleotides used in this study.

## References

1. Ending the Neglect to Attain the Sustainable Development Goals: A Road Map for Neglected Tropical Diseases 2021-2030. 1st ed. Geneva: World Health Organization; 2021.

2. Gull K. The cytoskeleton of trypanosomatid parasites. Annu Rev Microbiol. 1999;53: 629– 655. doi:10.1146/annurev.micro.53.1.629

3. Sinclair AN, De Graffenried CL. More than Microtubules: The Structure and Function of the Subpellicular Array in Trypanosomatids. Trends in Parasitology. 2019;35: 760–777. doi:10.1016/j.pt.2019.07.008

4. Janke C, Magiera MM. The tubulin code and its role in controlling microtubule properties and functions. Nat Rev Mol Cell Biol. 2020;21: 307–326. doi:10.1038/s41580-020-0214-3

5. Jackson AP, Vaughan S, Gull K. Evolution of tubulin gene arrays in Trypanosomatid parasites: genomic restructuring in Leishmania. BMC Genomics. 2006;7: 261. doi:10.1186/1471-2164-7-261

6. Nieuwenhuis J, Adamopoulos A, Bleijerveld OB, Mazouzi A, Stickel E, Celie P, et al. Vasohibins encode tubulin detyrosinating activity. Science. 2017;358: 1453–1456. doi:10.1126/science.aao5676

7. Aillaud C, Bosc C, Peris L, Bosson A, Heemeryck P, Van Dijk J, et al. Vasohibins/SVBP are tubulin carboxypeptidases (TCPs) that regulate neuron differentiation. Science. 2017;358: 1448–1453. doi:10.1126/science.aao4165

8. Nicot S, Gillard G, Impheng H, Joachimiak E, Urbach S, Mochizuki K, et al. A family of carboxypeptidases catalyzing α- and β-tubulin tail processing and deglutamylation. Sci Adv. 2023;9: eadi7838. doi:10.1126/sciadv.adi7838

9. Landskron L, Bak J, Adamopoulos A, Kaplani K, Moraiti M, Van Den Hengel LG, et al. Posttranslational modification of microtubules by the MATCAP detyrosinase. Science. 2022;376: eabn6020. doi:10.1126/science.abn6020

10. Raybin D, Flavin M. Enzyme which specifically adds tyrosine to the alpha chain of tubulin. Biochemistry. 1977;16: 2189–2194. doi:10.1021/bi00629a023

11. Barisic M, Silva E Sousa R, Tripathy SK, Magiera MM, Zaytsev AV, Pereira AL, et al. Microtubule detyrosination guides chromosomes during mitosis. Science. 2015;348: 799– 803. doi:10.1126/science.aaa5175

12. Robison P, Caporizzo MA, Ahmadzadeh H, Bogush AI, Chen CY, Margulies KB, et al. Detyrosinated microtubules buckle and bear load in contracting cardiomyocytes. Science. 2016;352: aaf0659. doi:10.1126/science.aaf0659

13. Sanyal C, Pietsch N, Ramirez Rios S, Peris L, Carrier L, Moutin M-J. The detyrosination/re-tyrosination cycle of tubulin and its role and dysfunction in neurons and cardiomyocytes. Seminars in Cell & Developmental Biology. 2023;137: 46–62. doi:10.1016/j.semcdb.2021.12.006

14. Konishi Y, Setou M. Tubulin tyrosination navigates the kinesin-1 motor domain to axons. Nat Neurosci. 2009;12: 559–567. doi:10.1038/nn.2314

15. Peris L, Thery M, Fauré J, Saoudi Y, Lafanechère L, Chilton JK, et al. Tubulin tyrosination is a major factor affecting the recruitment of CAP-Gly proteins at microtubule plus ends. The Journal of Cell Biology. 2006;174: 839–849. doi:10.1083/jcb.200512058

16. Hotta T, McAlear TS, Yue Y, Higaki T, Haynes SE, Nesvizhskii AI, et al. EML2-S constitutes a new class of proteins that recognizes and regulates the dynamics of tyrosinated microtubules. Current Biology. 2022;32: 3898–3910.e14. doi:10.1016/j.cub.2022.07.027

17. Sirajuddin M, Rice LM, Vale RD. Regulation of microtubule motors by tubulin isotypes and post-translational modifications. Nat Cell Biol. 2014;16: 335–344. doi:10.1038/ncb2920

18. Girão H, Macário-Monteiro J, Figueiredo AC, Silva E Sousa R, Doria E, Demidov V, et al. α-tubulin detyrosination fine-tunes kinetochore-microtubule attachments. Nat Commun. 2024;15: 9720. doi:10.1038/s41467-024-54155-8

19. McKenney RJ, Huynh W, Vale RD, Sirajuddin M. Tyrosination of α-tubulin controls the initiation of processive dynein-dynactin motility. EMBO J. 2016;35: 1175–1185. doi:10.15252/embj.201593071

20. Chhatre A, Stepanek L, Nievergelt AP, Alvarez Viar G, Diez S, Pigino G. Tubulin tyrosination/detyrosination regulate the affinity and sorting of intraflagellar transport trains on axonemal microtubule doublets. Nat Commun. 2025;16: 1055. doi:10.1038/s41467-025-56098-0

21. He HK, Kubo S, Lv QH, Kage A, Ichikawa M. Dynein-2 is tuned for the A-tubule of the ciliary doublet through tubulin tyrosination. 2025. doi:10.1101/2025.01.27.635025

22. Peris L, Wagenbach M, Lafanechère L, Brocard J, Moore AT, Kozielski F, et al. Motor-dependent microtubule disassembly driven by tubulin tyrosination. Journal of Cell Biology. 2009;185: 1159–1166. doi:10.1083/jcb.200902142

23. Strassel C, Magiera MM, Dupuis A, Batzenschlager M, Hovasse A, Pleines I, et al. An essential role for α4A-tubulin in platelet biogenesis. Life Sci Alliance. 2019;2: e201900309. doi:10.26508/lsa.201900309

24. Kerr JP, Robison P, Shi G, Bogush AI, Kempema AM, Hexum JK, et al. Detyrosinated microtubules modulate mechanotransduction in heart and skeletal muscle. Nat Commun. 2015;6: 8526. doi:10.1038/ncomms9526

25. Schneider A, Plessmann U, Weber K. Subpellicular and flagellar microtubules of Trypanosoma brucei are extensively glutamylated. J Cell Sci. 1997;110 (Pt 4): 431–437. doi:10.1242/jcs.110.4.431

26. Sherwin T, Schneider A, Sasse R, Seebeck T, Gull K. Distinct Localization and Cell Cycle Dependence of COOH Terminally Tyrosinolated a-Tubulin in the Microtubules of Trypanosoma brucei brucei. 1987.

27. van der Laan S, Lévêque MF, Marcellin G, Vezenkov L, Lannay Y, Dubra G, et al. Evolutionary Divergence of Enzymatic Mechanisms for Tubulin Detyrosination. Cell Rep. 2019;29: 4159–4171.e6. doi:10.1016/j.celrep.2019.11.074

28. Thein M, Wunderlich H, Brehm L, Wagner S, Weiss M, Ersfeld K. Effects of microtubule (de)tyrosination on the morphology and motility of Trypanosoma brucei and cross-talk with polyglutamylation. Biol Open. 2025;14: bio062270. doi:10.1242/bio.062270

29. Corrales RM, Vincent J, Crobu L, Neish R, Nepal B, Espeut J, et al. Tubulin detyrosination shapes *Leishmania* cytoskeletal architecture and virulence. Proc Natl Acad Sci USA. 2025;122: e2415296122. doi:10.1073/pnas.2415296122

30. Nisavic M, Chaze T, Bonnefoy S, Janke C, Bastin P, Matondo M, et al. Mass Spectrometry Reveals Novel Features of Tubulin Polyglutamylation in the Flagellum of Trypanosoma brucei. J Proteome Res. 2025;24: 3979–3989. doi:10.1021/acs.jproteome.5c00107

31. Casanova M, De Monbrison F, Van Dijk J, Janke C, Pagès M, Bastien P. Characterisation of polyglutamylases in trypanosomatids. International Journal for Parasitology. 2015;45: 121–132. doi:10.1016/j.ijpara.2014.09.005

32. Jentzsch J, Sabri A, Speckner K, Lallinger-Kube G, Weiss M, Ersfeld K. Microtubule polyglutamylation is important for regulating cytoskeletal architecture and motility in *Trypanosoma brucei*. Journal of Cell Science. 2020;133: jcs248047. doi:10.1242/jcs.248047

33. Jentzsch J, Wunderlich H, Thein M, Bechthold J, Brehm L, Krauß SW, et al. Microtubule polyglutamylation is an essential regulator of cytoskeletal integrity in *Trypanosoma brucei*. Journal of Cell Science. 2024; jcs.261740. doi:10.1242/jcs.261740

34. Li L, Stoeckert CJ, Roos DS. OrthoMCL: Identification of Ortholog Groups for Eukaryotic Genomes. Genome Res. 2003;13: 2178–2189. doi:10.1101/gr.1224503

35. Huerta-Cepas J, Szklarczyk D, Heller D, Hernández-Plaza A, Forslund SK, Cook H, et al. eggNOG 5.0: a hierarchical, functionally and phylogenetically annotated orthology resource based on 5090 organisms and 2502 viruses. Nucleic Acids Research. 2019;47: D309–D314. doi:10.1093/nar/gky1085

36. Jackson AP, Otto TD, Aslett M, Armstrong SD, Bringaud F, Schlacht A, et al. Kinetoplastid Phylogenomics Reveals the Evolutionary Innovations Associated with the Origins of Parasitism. Curr Biol. 2016;26: 161–172. doi:10.1016/j.cub.2015.11.055

37. Sasse R, Gull K. Tubulin Post-Translational Modifications And The Construction Of Microtubular Organelles In *Trypanosoma Brucei*. Journal of Cell Science. 1988;90: 577– 589. doi:10.1242/jcs.90.4.577

38. Sheriff O, Lim L-F, He CY. Tracking the Biogenesis and Inheritance of Subpellicular Microtubule in *Trypanosoma brucei* with Inducible YFP-*α* -Tubulin. BioMed Research International. 2014;2014: 1–12. doi:10.1155/2014/893272

39. Kalichava A, Ochsenreiter T. Ultrastructure expansion microscopy in *Trypanosoma brucei*. Open Biol. 2021;11: 210132. doi:10.1098/rsob.210132

40. Andre J, Kerry L, Qi X, Hawkins E, Drižytė K, Ginger ML, et al. An Alternative Model for the Role of RP2 Protein in Flagellum Assembly in the African Trypanosome. Journal of Biological Chemistry. 2014;289: 464–475. doi:10.1074/jbc.M113.509521

41. Stephan A, Vaughan S, Shaw MK, Gull K, McKean PG. An Essential Quality Control Mechanism at the Eukaryotic Basal Body Prior to Intraflagellar Transport. Traffic. 2007;8: 1323–1330. doi:10.1111/j.1600-0854.2007.00611.x

42. Corinne A, Perrine H, Pauline M, Quentin O, Alexandra MC, Valérie P-M, et al. Streptococcus pyogenes Cas9 ribonucleoprotein delivery for efficient, rapid and marker-free gene editing in Trypanosoma and Leishmania. 2023. doi:10.1101/2023.10.25.563974

43. McKean PG. Coordination of cell cycle and cytokinesis in Trypanosoma brucei. Curr Opin Microbiol. 2003;6: 600–607. doi:10.1016/j.mib.2003.10.010

44. Bodakuntla S, Yuan X, Genova M, Gadadhar S, Leboucher S, Birling M-C, et al. Distinct roles of α- and β-tubulin polyglutamylation in controlling axonal transport and in neurodegeneration. EMBO J. 2022;41: e111373. doi:10.15252/embj.2022111373

45. Bertiaux E, Morga B, Blisnick T, Rotureau B, Bastin P. A Grow-and-Lock Model for the Control of Flagellum Length in Trypanosomes. Current Biology. 2018;28: 3802–3814.e3. doi:10.1016/j.cub.2018.10.031

46. Ralston KS, Lerner AG, Diener DR, Hill KL. Flagellar Motility Contributes to Cytokinesis in *Trypanosoma brucei* and Is Modulated by an Evolutionarily Conserved Dynein Regulatory System. Eukaryot Cell. 2006;5: 696–711. doi:10.1128/EC.5.4.696-711.2006

47. Hill KL. Biology and Mechanism of Trypanosome Cell Motility. Eukaryot Cell. 2003;2: 200– 208. doi:10.1128/EC.2.2.200-208.2003

48. Shaw S, Roditi I. The sweet and sour sides of trypanosome social motility. Trends Parasitol. 2023;39: 242–250. doi:10.1016/j.pt.2023.01.001

49. Warn RM, Harrison A, Planques V, Robert-Nicoud N, Wehland J. Distribution of microtubules containing post-translationally modified alpha-tubulin during Drosophila embryogenesis. Cell Motil Cytoskeleton. 1990;17: 34–45. doi:10.1002/cm.970170106

50. Xiao H, El Bissati K, Verdier-Pinard P, Burd B, Zhang H, Kim K, et al. Post-translational modifications to Toxoplasma gondii alpha- and beta-tubulins include novel C-terminal methylation. J Proteome Res. 2010;9: 359–372. doi:10.1021/pr900699a

51. Velle KB, Kennard AS, Trupinić M, Ivec A, Swafford AJM, Nolton E, et al. Naegleria’s mitotic spindles are built from unique tubulins and highlight core spindle features. Curr Biol. 2022;32: 1247–1261.e6. doi:10.1016/j.cub.2022.01.034

52. Lai EY, Remillard SP, Fulton C. A beta-tubulin gene of Naegleria encodes a carboxy-terminal tyrosine. Aromatic amino acids are conserved at carboxy termini. J Mol Biol. 1994;235: 377–388. doi:10.1016/s0022-2836(05)80045-2

53. Redeker V, Levilliers N, Vinolo E, Rossier J, Jaillard D, Burnette D, et al. Mutations of tubulin glycylation sites reveal cross-talk between the C termini of alpha- and beta-tubulin and affect the ciliary matrix in Tetrahymena. J Biol Chem. 2005;280: 596–606. doi:10.1074/jbc.M408324200

54. Chen J, Kholina E, Szyk A, Fedorov VA, Kovalenko I, Gudimchuk N, et al. α-tubulin tail modifications regulate microtubule stability through selective effector recruitment, not changes in intrinsic polymer dynamics. Developmental Cell. 2021;56: 2016–2028.e4. doi:10.1016/j.devcel.2021.05.005

55. Paturle L, Wehland J, Margolis RL, Job D. Complete separation of tyrosinated, detyrosinated, and nontyrosinatable brain tubulin subpopulations using affinity chromatography. Biochemistry. 1989;28: 2698–2704. doi:10.1021/bi00432a050

56. Araujo Alves A, Jung J, Moneron G, Vaucelle H, Fort C, Buisson J, et al. Intraflagellar transport selectivity occurs within the proximal portion of the trypanosome flagellum. J Cell Biol. 2025;224: e202501088. doi:10.1083/jcb.202501088

57. Mallet A, Bastin P. Restriction of intraflagellar transport to some microtubule doublets: An opportunity for cilia diversification? Bioessays. 2022;44: e2200031. doi:10.1002/bies.202200031

58. Shaw S, DeMarco SF, Rehmann R, Wenzler T, Florini F, Roditi I, et al. Flagellar cAMP signaling controls trypanosome progression through host tissues. Nat Commun. 2019;10: 803. doi:10.1038/s41467-019-08696-y

59. Ralston KS, Kabututu ZP, Melehani JH, Oberholzer M, Hill KL. The Trypanosoma brucei flagellum: moving parasites in new directions. Annu Rev Microbiol. 2009;63: 335–362. doi:10.1146/annurev.micro.091208.073353

60. Tashyreva D, Faktorová D, Horák A, Lukeš J, Archibald JM, Oatley G, et al. The genome sequences of the diplonemid protist Rhynchopus euleeides YPF1915 and its bacterial endosymbiont Candidatus Syngnamydia salmonis (Chlamydiota). Wellcome Open Res. 2025;10: 233. doi:10.12688/wellcomeopenres.24014.1

61. Zavadska D, Packer JA, Tashyreva D, Kamikawa R, Simpson A, Richter DJ. Phylogenomics of free-living neobodonids reveals they are a paraphyletic group from which all other metakinetoplastids are descended. Evolutionary Biology; 2025. doi:10.1101/2025.10.03.680260

62. Butenko A, Opperdoes FR, Flegontova O, Horák A, Hampl V, Keeling P, et al. Evolution of metabolic capabilities and molecular features of diplonemids, kinetoplastids, and euglenids. BMC Biol. 2020;18: 23. doi:10.1186/s12915-020-0754-1

63. Beneke T, Madden R, Makin L, Valli J, Sunter J, Gluenz E. A CRISPR Cas9 high-throughput genome editing toolkit for kinetoplastids. R Soc Open Sci. 2017;4: 170095. doi:10.1098/rsos.170095

64. Alves AA, Gabriel HB, Bezerra MJR, de Souza W, Vaughan S, Cunha-E-Silva NL, et al. Control of assembly of extra-axonemal structures: the paraflagellar rod of trypanosomes. J Cell Sci. 2020;133: jcs242271. doi:10.1242/jcs.242271

65. Louvel V, Haase R, Mercey O, Laporte MH, Eloy T, Baudrier É, et al. iU-ExM: nanoscopy of organelles and tissues with iterative ultrastructure expansion microscopy. Nat Commun. 2023;14: 7893. doi:10.1038/s41467-023-43582-8

66. Piperno G, LeDizet M, Chang XJ. Microtubules containing acetylated alpha-tubulin in mammalian cells in culture. J Cell Biol. 1987;104: 289–302. doi:10.1083/jcb.104.2.289

67. Imhof S, Knüsel S, Gunasekera K, Vu XL, Roditi I. Social Motility of African Trypanosomes Is a Property of a Distinct Life-Cycle Stage That Occurs Early in Tsetse Fly Transmission. Hill KL, editor. PLoS Pathog. 2014;10: e1004493. doi:10.1371/journal.ppat.1004493

